# Tissue-specific degradation of essential centrosome components reveals distinct microtubule populations at microtubule organizing centers

**DOI:** 10.1101/363325

**Authors:** Maria D. Sallee, Jennifer C. Zonka, Taylor D. Skokan, Brian C. Raftrey, Jessica L. Feldman

## Abstract

Non-centrosomal microtubule organizing centers (ncMTOCs) are found in most differentiated cells, but how these structures regulate microtubule organization and dynamics is largely unknown. We optimized a tissue-specific degradation system to test the role of the essential centrosomal microtubule nucleators γ-tubulin ring complex (γ-TURC) and AIR-1/Aurora A at the apical ncMTOC, where they both localize in *C. elegans* embryonic intestinal epithelial cells. As at the centrosome, the core γ-TURC component GIP-1/GCP3 is required to recruit other γ-TuRC components to the apical ncMTOC including MZT-1/MZT1, characterized here for the first time in animal development. In contrast, AIR-1 and MZT-1 were specifically required to recruit γ-TuRC to the centrosome, but not to centrioles or to the apical ncMTOC. Surprisingly, microtubules remain robustly organized at the apical ncMTOC upon γ-TuRC and AIR-1 co-depletion, and upon depletion of other known microtubule regulators including TPXL-1/TPX2, ZYG-9/chTOG, PTRN-1/CAMSAP, and NOCA-1/Ninein. However, loss of GIP-1 removed a subset of dynamic EBP-2/EB1-marked microtubules, and the remaining dynamic microtubules grew faster. Together, these results suggest that different MTOCs use discrete proteins for their function, and that the apical ncMTOC is composed of distinct populations of γ-TuRC-dependent and independent microtubules that compete for a limited pool of resources.

## Introduction

Described nearly 50 years ago, microtubule organizing centers (MTOCs) generate specific spatial patterns of microtubules as needed for cell function [1]. The best-studied MTOC is the centrosome, a non-membrane bound organelle that organizes microtubules into a radial array from its pericentriolar material (PCM) or from subdistal appendages attached to the mother centriole. However, in many types of differentiated cells, microtubules are organized at non-centrosomal sites to accommodate diverse cell functions. In animal cells, these non-centrosomal MTOCs (ncMTOCs) can be found in the axons and dendrites of neurons, around the nuclear envelope of skeletal muscle cells, at the apical surface of epithelial cells, and at the Golgi complex [2–8]. ncMTOCs can promote non-radial arrangements of microtubules, such as the linear arrays of microtubules present along the apico-basal axis in epithelial cells. How ncMTOCs are established and whether they are composed of the same proteins that impart MTOC activity at the centrosome is largely unknown.

In general, MTOCs can be defined as cellular sites that nucleate, anchor, and stabilize microtubules; however, the molecular basis for these functions has been elusive [9,10]. Because of the inherent structural and chemical polarity of microtubule polymers, microtubules are nucleated and anchored at their minus ends. Thus, a defining feature of MTOCs is that they interact with microtubule minus ends. The first microtubule minus-end protein described was γ-tubulin, which together with GCP2 and GCP3, forms the γ-tubulin small complex (γ-TuSC) [11]. In some organisms, additional γ-tubulin complex proteins (GCPs) combine with the γ-TuSC to form the larger γ-tubulin ring complex (γ-TURC); organisms lacking these additional GCPs are thought to oligomerize γ-TuSCs into similar ring complexes [12,13]. To date, only the γ-TuSC components TBG-1/γ-tubulin, GIP-1/GCP3, and GIP-2/GCP2 have been identified in *C. elegans*, suggesting that *C. elegans* γ-TURC may share the yeast γ-TURC composition. As our experiments do not distinguish between γ-TUSC and γ-TuRC, we will use the term γ-TURC for simplicity. A putative *C. elegans* ortholog of MOZART1 (mitotic spindle-organizing protein associated with a ring of γ-tubulin, MZT1), a γ-TURC-interacting protein and proposed γ-TURC component, was identified based on sequence homology to the *Arabidopsis* ortholog, but its function has not been investigated [14]. γ-TURC has microtubule nucleation capacity and can also cap the minus ends of microtubules, preventing minus-end growth or depolymerization [15,16]. Whether γ-TURC predominantly functions as a nucleator or as a minus-end cap or anchor *in vivo* is a matter of debate.

Although γ-TURC is essential in organisms ranging from yeast to humans, γ-TuRC depletion does not result in the elimination of all microtubules from the cell, suggesting that other mechanisms exist to grow and anchor microtubules at MTOCs. γ-TURC removal *in vivo* has severely deleterious effects on mitosis, but microtubules are still present [17–19]. The presence of microtubules in dividing *C. elegans* embryonic cells appears to rely on both γ-TURC function and the mitotic kinase AIR-1/Aurora A [20,21], as only depletion of both TBG-1 and AIR-1 from dividing cells results in the elimination of centrosomal microtubules. Whether γ-TuRC and AIR-1, or other essential centrosomal MTOC proteins, function redundantly to build microtubules at ncMTOCs in animal cells is unknown, most notably because of the early requirement of these proteins in mitosis that prohibits an assessment of any later roles during differentiation.

In *C. elegans* embryonic intestinal cells, MTOC function is reassigned from the centrosome during mitosis to the apical surface as cells begin to polarize [6], thereby establishing an apical ncMTOC in each cell. Intestinal cells all derive from the ‘E’ blastomere, undergoing 4 rounds of division before polarizing at the ‘E16’ stage when the intestinal primordium is comprised of 16 epithelial cells (S1A Fig, [22]). Shortly after the E8-E16 division, the E16 cells follow a stereotypical pattern of polarization and establish their apical surfaces facing a common midline [6], the eventual site of the lumen of the epithelial tube. ncMTOCs positioned along these apical surfaces nucleate and organize microtubules into fountain-like arrays emanating away from the midline on either side [6,23,24]. Intriguingly, many centrosomal MTOC proteins including AIR-1 and GIP-1 also localize to this ncMTOC [6]. Because the earliest stages of ncMTOC formation can be easily tracked, the embryonic intestinal primordium provides an ideal system in which to test the role of specific proteins in ncMTOC establishment *in vivo.*

Here, we test the hypothesis that the proteins required to build microtubules at the centrosome play a similar role at ncMTOCs. To do this, we optimized an existing tissue-specific degradation strategy to test the role of GIP-1 and AIR-1 at ncMTOCs in *C. elegans* embryonic intestinal cells. As at the centrosome, we find that GIP-1 is required to localize the other γ-TURC members, TBG-1 and GIP-2. Additionally, we showed that a predicted ortholog of the γ-TuRC protein MZT1 is essential in *C. elegans* and colocalizes with γ-TURC in all contexts but is uniquely required for localization of γ-TURC to the PCM, and not to the centriole or to the apical ncMTOC. This differential requirement for proteins at the centrosome versus the apical ncMTOC was a common trend, as AIR-1 was also only required to localize GIP-1 and TAC-1 to the PCM, but not to the apical ncMTOC. In addition to GIP-1 and AIR-1, we assessed the requirement of other known microtubule regulators, including ZYG-9/chTOG, PTRN-1/CAMSAP, NOCA-1/Ninein, and TPXL-1/TPX2. Surprisingly, we found that overall the depletion of these proteins did not disrupt apical microtubule organization. Furthermore, removal of GIP-1 had only a minor effect on microtubule dynamics at the apical ncMTOC; a subset of microtubules was perturbed as indicated by a change in EBP-2/EB1 localization and dynamics. These results highlight the differences between the centrosome and other MTOCs and suggest that ncMTOCs are composed of at least two populations of microtubules, γ-TuRC-dependent and γ-TuRC-independent.

## Results

### An optimized ZF/ZIF-1 degradation system allows for tissue-specific degradation of early essential proteins

To test the role of γ-TURC and AIR-1 at ncMTOCs, we needed a strategy to deplete proteins essential in the early embryo (early essential proteins) at later stages of development. For example, we had previously been unable to assess the function of γ-TURC components in differentiated cells *in vivo* as their depletion causes severe mitotic defects that result in early embryonic lethality. Tissue-specific degradation strategies have provided a means to deplete such early essential proteins [25,26]. We therefore optimized an existing tissue-specific degradation system (Fig 1A, B). The germline cell fate determinant PIE-1 is degraded in somatic cells in the early embryo ([27], Fig 1C). This degradation requires a zinc finger domain 1 (ZF) on PIE-1 and the SOCS-box protein ZIF-1, which targets PIE-1 for degradation by an E3 ubiquitin ligase [27]. Previous reports found that the ZF domain could be added to any protein of interest and that protein is degraded in somatic cells by ZIF-1 ([28], Fig 1E, G). However, endogenous ZIF-1 is only expressed in the early embryo so degradation of targets later in development requires exogenous expression of ZIF-1 [29]. The major drawback of this system is that degradation of early essential proteins by endogenous ZIF-1 leads to an early arrest. We found that a *zif*-*1* deletion mutant is viable (92% ± 8.4% embryonic viability in *zif*-*1(gk117)* worms compared to 99% ± 1.9% in N2 worms) despite the apparent loss of ZIF-1 activity (Fig 1C, D). Using a *zif*-*1* mutant background (‘*zif*-*1(*-)’), we can tag any gene with the ZF domain using CRISPR/Cas9 and the resulting ZF-tagged protein is not degraded (Fig 1F, H, I, L, O), thus allowing for normal development. We then express ZIF-1 under the control of a tissue-specific promoter to degrade ZF-tagged targets.

**Fig. 1.**
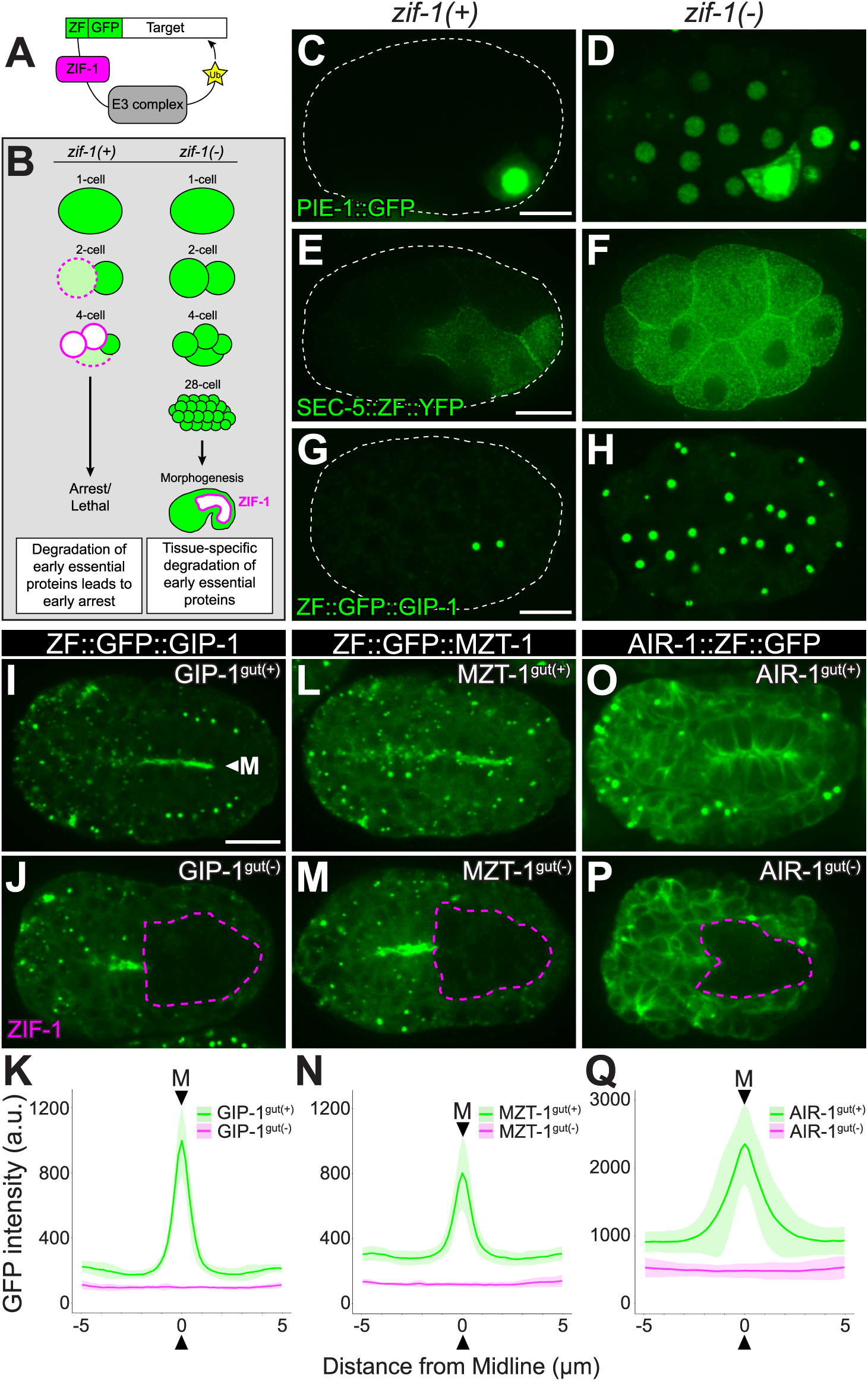
A tissue-specific degradation system to deplete early essential proteins A, B) Cartoons depicting tissue-specific protein degradation scheme (adapted from [29]). In the presence of endogenous ZIF-1 (*zif-1(+)*), ZF-tagged targets are degraded in somatic cells leading to an early arrest in the case of early essential proteins. *zif*-*1(*-) mutants (*zif*-*1(gk117)*) fail to degrade endogenous ZF-tagged targets. C, E, G) PIE-1::GFP, a natural target of ZIF-1, and SEC-5::ZF::GFP and ZF::GFP::GIP-1, two heterologous ZIF-1 targets, are degraded in somatic cells, but not in the germ cell precursors. Dashed white lines outline embryos. D, F, H) In *zif*-*1(*-) embryos, ZF-containing targets are not degraded. I, L, O) Bean-stage *zif*-*1(*-) mutants in which indicated endogenous loci have been tagged with ZF::GFP but ZIF-1 is not expressed (‘gut(+)’). Note the localization to the apical surfaces of intestinal cells (‘M’, arrowhead). J, M, P) Expression of ZIF-1 in intestinal cells (magenta dashed lines) results in tissue-specific degradation (‘gut(-)’). Scale bar is 10 μm. K, N, Q) Average 10 μm line intensity profiles across the apical midline (‘M’, arrowhead) of control embryos (green lines show mean, lighter green shading indicate standard deviation) show a peak in GFP signal intensity in GIP-1^gut(+)^ (n = 12), MZT-1^gut(+)^ (n = 11), and AIR-1^gut(+)^ (n = 13) embryos, and no peak and reduced cytoplasmic signal upon degradation of tagged proteins (magenta lines show mean, lighter magenta shading indicate standard deviation) in GIP-1^gut(-)^ (n = 12), MZT-1^gut(-)^ (n = 10), and AIR-1^gut(-)^ (n = 15) embryos. See S1 Fig, S2 Fig, and S3 Fig for details.

Using this strategy, we tagged the γ-TURC component GIP-1/GCP3, the predicted MZT1 ortholog W03G9.8 which we hereafter refer to as MZT-1 (see below), and the mitotic kinase AIR-1/Aurora A with ZF::GFP, allowing us to monitor protein expression and localization and to degrade each protein with exogenous ZIF-1 expression (Fig 1I-Q). As expected, GIP-1 localized to the apical ncMTOC in intestinal cells and AIR-1 decorated microtubules (Fig 1I, O, [6]). ZIF-1 was then expressed using the promoter for the *elt*-*2* gene, which is expressed exclusively in the intestine starting at intestinal stage E2 (S1A Fig). ZIF-1 expression led to intestine-specific removal of GIP-1, MZT-1, or AIR-1 (‘GIP-1^gut(-)^’, ‘MZT-1^gut(-)^’, ‘AIR-1^gut(-)^’) as demonstrated by the loss of both apical and cytoplasmic GFP signal (Fig 1I-Q, S1B-E Fig, S2A-D’ Fig). We quantified this intestine-specific depletion in two ways. First, we measured the total amount of reduction of GFP signal in the intestinal primordium of ‘gut(-)’ embryos as compared to ‘gut(+)’ siblings that lacked the ZIF-1-expressing array (% GFP depletion: GIP-1^gut(-)^ 93.1%, MZT-1^gut(-)^ 92.1%, AIR-1^gut(-)^ 82.1%, S3A Fig). This is likely an underestimate, due to the out of focus light contributed by non-degraded ZF::GFP in non-intestinal cells that complicates this assessment, especially when highly expressed genes like *air*-*1* were tagged. We also took line scans across the midline of the intestinal primordium in both gut(-) and gut(+) embryos to measure apical enrichment (Fig 1K, N, Q, S3B, D Fig). We see no significant apical enrichment of GFP signal in GIP-1^gut(-)^, MZT-1^gut(-)^, AIR-1^gut(-)^ embryos, and a significant reduction in both apical and cytoplasmic GFP intensity as compared to gut(+) control embryos. This degradation strategy can also be used to co-deplete AIR-1 and GIP-1 ([GIP-1;AIR-1]^gut(-)^, S1D Fig, S2A-D’ Fig, Materials and Methods). Thus, the *zif*-*1* mutant coupled with the ZIF-1/ZF degradation system provides an effective tool for depleting early essential proteins in a tissue-specific manner.

### Degradation of GIP-1/GCP3, MZT-1, and/or AIR-1/Aurora A in intestinal cells leads to mitotic defects, but does not impair intestinal differentiation

To test the role of γ-TURC and AIR-1 in establishing the apical ncMTOC, we needed to effectively remove these proteins from intestinal cells prior to the E16 stage when cells reassign MTOC function to the apical membrane. Intestinal differentiation in *C. elegans* proceeds in the absence of cell division [30], suggesting that mitotic defects in intestinal cells per se would not affect their ability to build an ncMTOC. Thus, we could begin degradation of the desired targets during the intestinal divisions to ensure they would be effectively cleared by the time cells began to polarize and establish the apical ncMTOC.

In *C. elegans*, loss of maternal AIR-1 results in severe mitotic defects including multinucleate cells, polyploidy, disorganized microtubules, and failed centrosome separation [31,32]; the additional removal of γ-TURC components results in monopolar spindles and loss of centrosomal microtubules in the first cell division [20,21]. We thus used mitotic defects in intestinal cells as a phenotypic read out for effective degradation of GIP-1 and AIR-1 prior to polarization. The E blastomere undergoes 4 rounds of division to generate the polarized 16-cell primordium (E16, Fig 2A). As ZIF-1 was expressed from an early promoter (*elt-2*p) that is active beginning around E2-E4, we expected that successful removal of γ-TURC and AIR-1 should result in polarized intestinal primordia with between 2 and 16 cells. Indeed, we found that degradation of GIP-1, MZT-1, or AIR-1 resulted in embryos with 8.6 ± 2.3, 7.6 ± 1.8, or 9.2 ± 1.5 intestinal nuclei, respectively, and that degradation of both AIR-1 and GIP-1 resulted in embryos with 4.0±0.0 intestinal nuclei (Fig 2A, B). Expression of ZIF-1 from a promoter that is active around E8 (*ifb*-*2*p) also significantly reduced the number of intestinal nuclei, although to a lesser extent (Fig 2A, B, S1A Fig). As further proof that we were effectively depleting the desired targets, we found that embryos with decreased or no zygotic *air*-*1*, and with only a maternal supply of ZF::GFP-tagged AIR-1 (‘AIR-1^*^’), had intestinal nuclear numbers indistinguishable from AIR-1^gut(-)^ embryos (see Materials and Methods, 9.1 ± 2.2, Fig 2B). We frequently observed mitotic defects in AIR-1^gut(-)^ and GIP-1^gut(-)^ embryos, such as scattered condensed chromosomes, binucleate cells, and abnormal mitotic spindles (S2I Fig, S1 Movie). These results are consistent with the reported role for γ-TURC and AIR-1 in mitosis and suggest that GIP-1 and AIR-1 are effectively depleted from intestinal cells beginning at approximately E4.

**Fig. 2.**
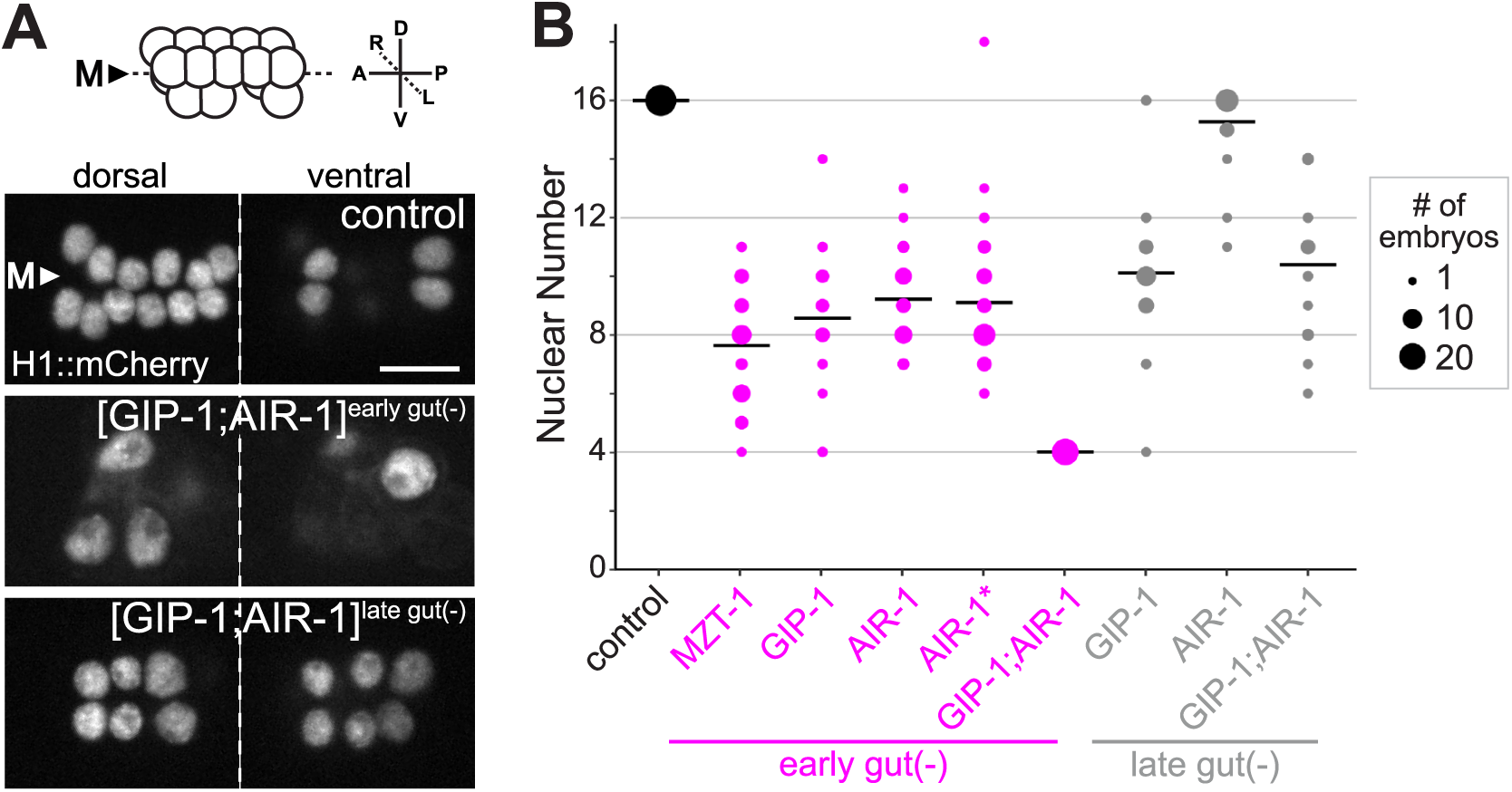
Degradation of γ-TuRC components and/or AIR-1 results in intestinal nuclear number defects A) Top: Cartoon depicting the organization of a wild-type bean-stage embryonic intestinal primordium, showing the dorsal and ventral tiers of cells. Below: Bean-stage embryos of indicated genotypes expressing intestine-specific histone::mCherry. Nuclei are organized into a dorsal (left) and ventral (right) tier of cells. Scale bar is 5 μm. B) Expression of ZIF-1 from an early intestinal promoter (*elt-2*p, ‘early gut(-)’, magenta dots) or late intestinal promoter (*ifb*-*2*p, ‘late gut(-)’, gray dots) to degrade MZT-1, GIP-1, AIR-1, or GIP-1;AIR-1 perturbs intestinal nuclear number as compared to control embryos (black dots), which have ZF::GFP tagged GIP-1 and AIR-1, but lack *elt*-*2*p::zif-1. ‘AIR-1^*^’ embryos are from *air*-*1(0/*[AIR-1::ZF::GFP]) mothers, and carry zero, one, or two copies of *air*-*1(0).* The distribution of nuclear number in AIR-1 and AIR-1^*^ embryos are not significantly different (two-tailed t-test, p = 0.81). Dot size indicates number of embryos. Early gut(-): control n = 28, MZT-1 n = 38, GIP-1 n = 21, AIR-1 n = 28, AIR-1^*^ n = 39, GIP-1;AIR-1 n = 20. Late gut(-): GIP-1 n = 28, AIR-1 n = 22, GIP-1;AIR-1 n = 18.

We next tested whether intestinal cells can polarize and differentiate in the absence of GIP-1 or AIR-1. In intestinal cells depleted of both GIP-1 and AIR-1, the apical polarity protein PAR-3, whose localization to the apical surface is a hallmark of apico-basal polarity [33], was unperturbed (S2E-H Fig). Similarly, we observed intestine-specific lysosome-like organelles known as gut granules (Fig 3D), which are hallmarks of intestinal differentiation [23,30]. Together, these results confirmed that we could use tissue-specific degradation to deplete GIP-1 and AIR-1 prior to apical ncMTOC formation without any dramatic effects on intestinal differentiation.

**Fig. 3.**
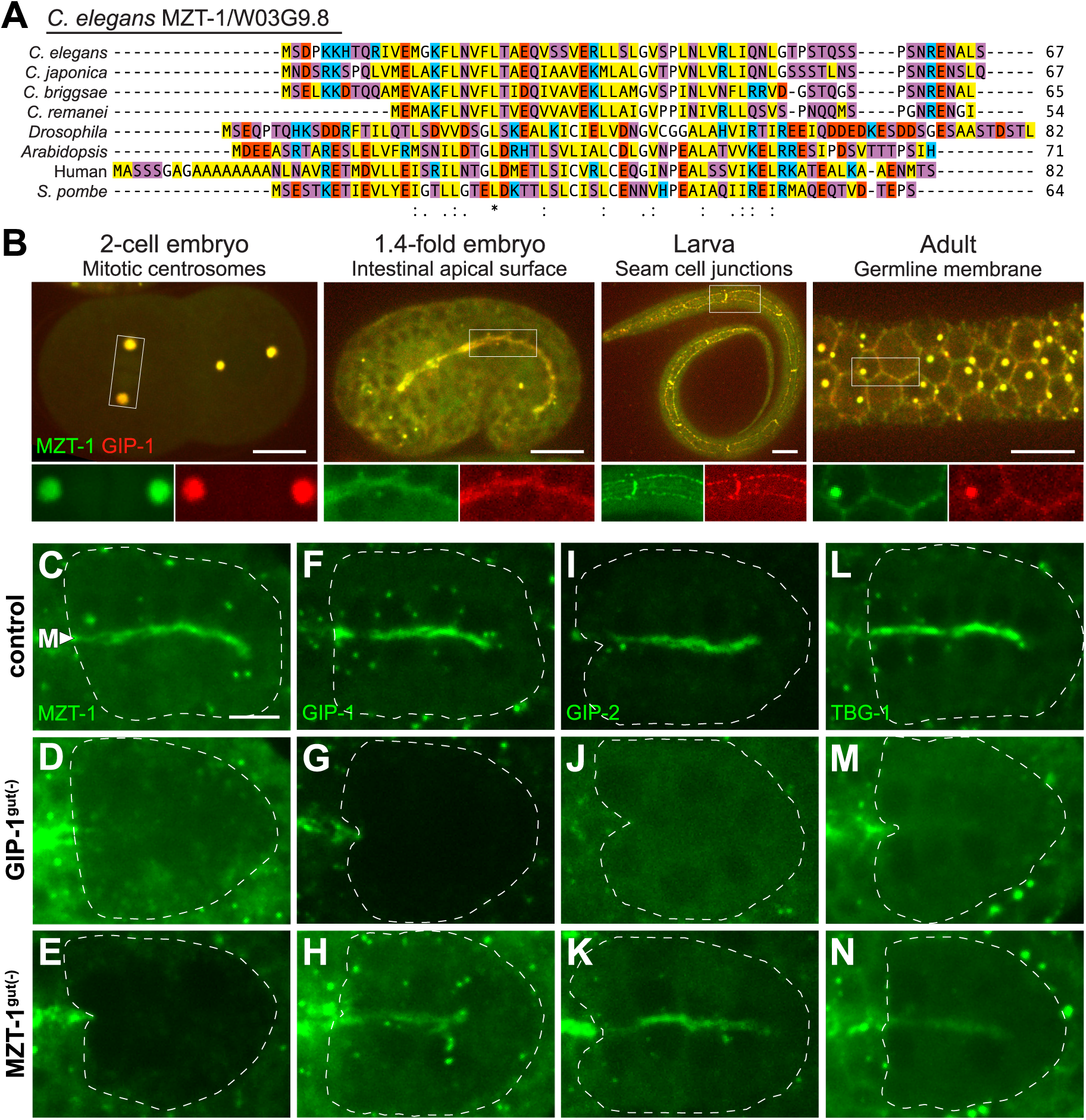
W03G9.8 is a MZT1 ortholog but, unlike GIP-1, is not required forγ-TuRC localization to the apical ncMTOC A) Alignment of W03G9.8 (renamed MZT-1) with *Caenorhabditis*, *Drosophila*, *Arabidopsis*, human and *S. pombe* orthologs. Sequences are also in S1 Text. B) Projected optical sections (embryos and germline) or a single optical section (seam cells) showing localization of endogenous tagRFP::GIP-1 and ZF::GFP::MZT-1 at centrosomes in the 2-cell embryo, the apical ncMTOC of the pharynx and intestinal primordia in a 1.4-fold embryo, the cell junctions of larval seam cells, and the membrane and centrosomes of the adult germline. Scale bar is 10 μm, insets are 2x magnifications with channels shown separately. C-N) Images are projected optical sections through the midline of live bean-stage embryos. The intestinal primordium is outlined by white dashed lines. Embryos are expressing a GFP-tagged version of the indicated protein from either the endogenous locus (GFP::MZT-1 in C, D; GFP::GIP-1 in F, H), an extrachromosomal array (GFP::GIP-2 in I-K), or a maternally expressed single copy insertion (TBG-1::GFP in L-N). Control *zif*-*1(*-) embryos expressing *elt*-*2*p::*zif*-*1* but lacking ZF-tagged CRISPR alleles are shown in (C, F, I, L), GIP-1^gut(-)^ embryos in (D, G, J, M) and MZT-1^gut(-)^ embryos in (E, H, K, N). Apical GFP::MZT-1 was observed in 29/29 control and in 0/29 GIP-1^gut(-)^ embryos; apical GFP::GIP-1 was observed in 17/17 control and 28/28 MZT-1^gut(-)^ embryos; apical GFP::GIP-2 was observed in 37/37 control, 1/24 GIP-1^gut(-)^, and 19/21 MZT-1^gut(-)^ embryos; and apical GFP::TBG-1 was observed in 19/19 control, 3/16 GIP-1^gut(-)^, and 14/15 MZT-1^gut(-)^ embryos. ZF::GFP::MZT-1 depletion is shown in (E) and ZF::GFP::GIP-1 depletion is shown in (G), and gut granules are visible in (D). Note that in GIP-1^gut(-)^ and MZT-1^gut(-)^ embryos, the non-degraded ZF::GFP-tagged protein is still present outside of the intestinal primordium. Scale bar is 5 μm.

### *W03G9.8*, the predicted C. elegans MZT1 ortholog, is an essential gene that colocalizes with GIP-1

MZT proteins have been shown to recruit γ-TURC to spindle poles in plants, fungi, and human cell culture lines, but have not been characterized *in vivo* in animal cells [14,34–38]. A previous study of the *Arabidopsis* homolog of the small protein MZT1 (GIP1) identified the uncharacterized gene *W03G9.8* (hereafter called *mzt*-*1*) by sequence homology as a potential MZT1 ortholog in *C. elegans* (Fig 3A, [14]). We tagged the endogenous locus of *mzt*-*1* with ZF::GFP to monitor endogenous MZT-1 localization and assess its function. In all embryonic, larval, and adult tissues examined, we observed consistent colocalization of ZF::GFP::MZT-1 with tagRFP::GIP-1, including at centrosomes in the early embryo, the apical membrane of the intestinal and pharynx primordia, the cell junctions of seam cells, and the plasma membrane and centrioles of the germline (Fig 3B). In the intestinal primordium, MZT-1 localized to centrioles and the apical ncMTOC in an identical pattern to all three γ-TURC components GIP-1/GCP3, GIP-2/GCP2 and TBG-1/γ-tubulin (Fig 3C, F, I, L). Like other γ-TURC members *tbg*-*1* and *gip*-*1* [17,39], we find that *mzt*-*1* is also required for embryonic viability. In a *zif*-*1(*+) background, ZF::GFP::MZT-1 is degraded ubiquitously in somatic cells by endogenous ZIF-1, and we observed fully penetrant maternal effect lethality (0/420 embryos hatched). By contrast, in control embryos (*zif*-*1(*+); GFP::MZT-1), endogenous GFP::MZT-1 is not degraded by ZIF-1 and the embryos are viable (340/344 hatched and grew to adulthood). Based on sequence homology, colocalization with GIP-1, and embryonic lethality, our results suggest that *W03G9.8/mzt*-*1* is the *C. elegans* ortholog of MZT1.

### GIP-1/GCP3 is required to localize other γ-TURC components to the apical ncMTOC

Our ability to deplete GIP-1 (Fig 3G) and MZT-1 (Fig 3E) in intestinal cells afforded us the opportunity to test their role in the recruitment of each other and of other γ-TURC components to the apical ncMTOC. At the centrosome, γ-TURC components exhibit interdependent localization [17]. We see a similar requirement for GIP-1 in localizing MZT-1, GIP-2, and TBG-1 to the apical membrane (compare Fig 3C, I, L to 3D, J, M), suggesting that as at the centrosome, these proteins localize to the apical ncMTOC as a complex. Additionally, the disrupted localization of other γ-TURC components upon GIP-1 depletion further confirmed that we had successfully perturbed GIP-1 function. In contrast to GIP-1^gut(-)^ embryos, GIP-1, GIP-2, and TBG-1 all localized apically in MZT-1^gut(-)^ embryos (Fig 3H, K, N), suggesting that MZT-1 is not required to recruit γ-TuRC to the apical ncMTOC.

### MZT-1 is required for recruitment of γ-TURC to the PCM, but not to the centrioles or apical ncMTOC

The surprising finding that γ-TURC still localizes to the apical ncMTOC upon MZT-1 depletion led us to further investigate the role of MZT-1 in γ-TURC recruitment to different sites. γ-TURC localizes to three distinct locations around the time of polarization. During the divisions that precede intestinal polarization, γ-TuRC accumulation at the PCM is coupled with the cell cycle. GIP-1 is recruited to the PCM of the centrosome as cells enter mitosis (Fig 4A). Following mitosis, the centrosome migrates to the lateral membrane and the PCM is greatly reduced. In this lateral configuration, we frequently observe two γ-TuRC positive puncta per cell, which by electron microscopy are centrioles associated with a small shell of PCM (‘paired centrosomes’, [6], Fig 4B, ‘interphase’ inset, blue double arrows). The paired centrosomes then migrate to the apical surface, where they appear as naked centrioles that completely lack PCM by electron microscopy [6]. At this stage, GIP-1 localizes to both the centrioles and the apical ncMTOC (Fig 4C). By contrast, MZT-1^gut(-)^ embryos localized GIP-1 to their centrioles, but failed to recruit GIP-1 to the PCM during mitosis (compare Fig 4A to 4D). We frequently saw inappropriate numbers of centrioles in MZT-1^gut(-)^ embryos (number of GIP-1 positive foci in control: 1.8 ± 0.5, n = 64 cells, MZT-1^gut(-)^: 3.0 ± 1.0, n = 134 cells, two-tailed t-test: p = 4.4 × 10^−24^), which may be an indirect consequence of earlier mitotic defects and which appeared to result in multipolar spindles (Fig 4D). Following mitosis in MZT-1^gut(-)^ embryos, GIP-1 remained associated with centrioles and accumulated at the apical ncMTOC as in control embryos (Fig 3H, 4B, C, E, F). To our knowledge, these results provide the first *in vivo* role for MZT-1/MZT1 in animal cells, suggesting that MZT-1 is a PCM-specific linker for γ-TURC in dividing cells. Consistent with these findings, human MZT1 appears to promote the targeting and activation of an intact γ-TuRC to the centrosome in human tissue culture cells [38]. In *C. elegans*, MZT-1 localization tracks with γ-TURC localization at the centriole, the PCM, and the apical ncMTOC. However, MZT-1 is not required to target γ-TURC to the apical ncMTOC but requires GIP-1 to localize there, suggesting that MZT-1 is stably associated with the complex even when not playing a targeting role.

**Fig. 4.**
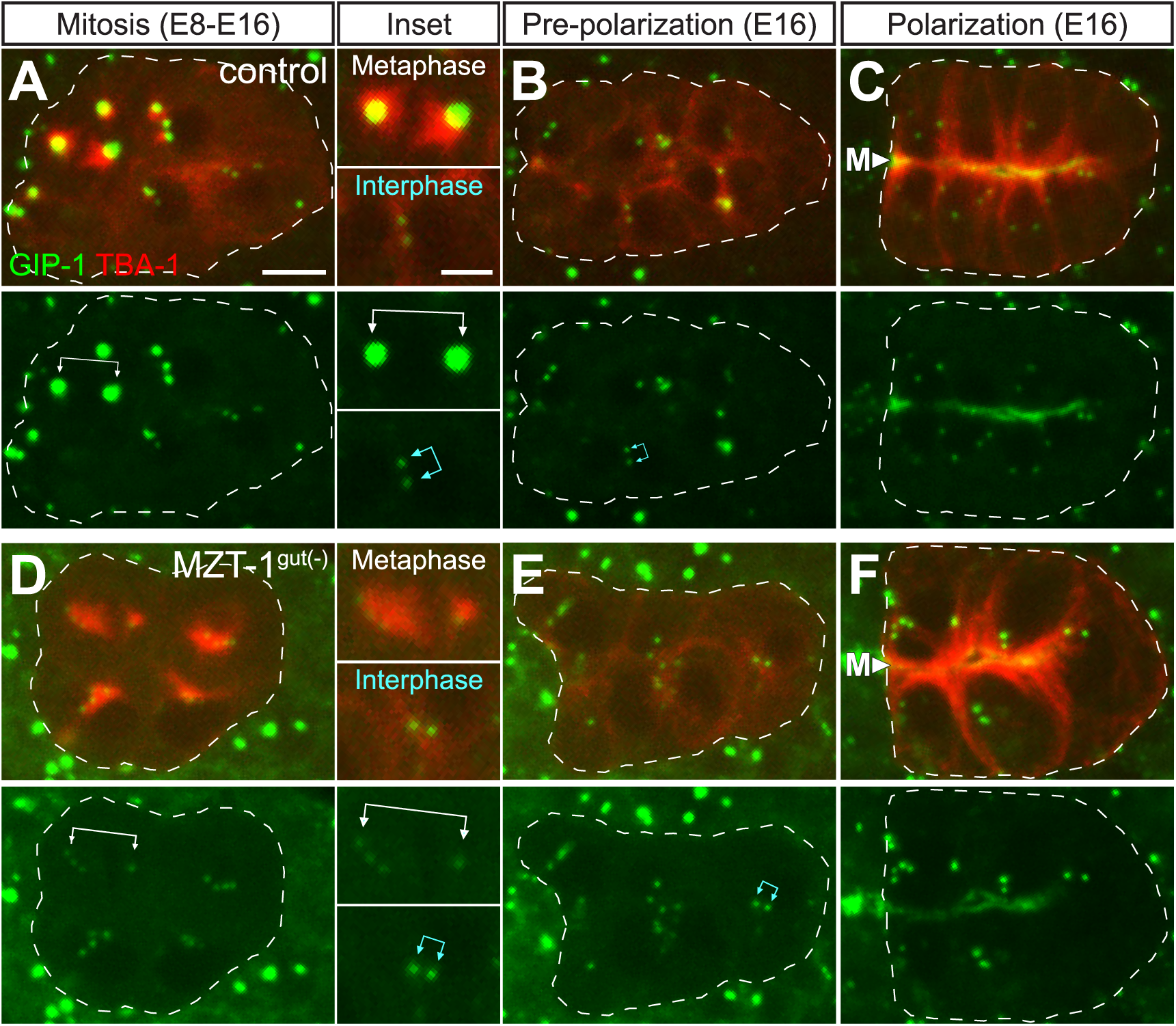
MZT-1 is required to localize GIP-1 to the centrosome, but not to the centriole or apical ncMTOC Images are projected optical sections through the intestinal primordium of live *zif*-*1(*-) control (A-C) and MZT-1^gut(-)^ (D-F) embryos as they develop from the E8-E16 divisions (A, D) through polarization (B, C, and E, F). The intestinal primordium is outlined by white dashed lines. Embryos express endogenous GFP::GIP-1 (green, single channel shown below each panel) and intestine-specific mCherry::TBA-1/α-tubulin (red). In addition, MZT-1^gut(-)^ embryos have nondegraded ZF::GFP::MZT-1 in non-intestinal cells. Note that the divisions in A are not completely synchronous; anterior (leftmost) cells are in mitosis, while more posterior (rightmost) cells have not yet started to divide. Top inset to the right of A and D shows a metaphase spindle. *zif*-*1(*-) control embryos (n = 24 centrosomes) recruit a large amount of GIP-1 to the PCM as compared to MZT-1^gut(-)^ embryos (n = 18 centrosomes, white joined arrows). MZT-1^gut(-)^ intestinal cells have abnormal numbers of centrioles and multipolar spindles. (B, E) After division, centrosomes shed their PCM (blue joined arrows in bottom inset at left). C, F) In polarized intestinal primordia, GIP-1 localizes to the apical surfaces at the midline (‘M’). Centriole and apical localization of GIP-1 appears unaffected in MZT-1^gut(-)^ embryos (n = 28/28). Scale bar is 5 μm or 2.5 μm in insets.

### γ-TuRC components are not required to recruit other minus-end proteins to the apical ncMTOC

Many microtubule minus-end proteins are found at MTOCs, reflecting the defining ability of MTOCs to nucleate and stabilize microtubule minus ends. To date, only a handful of these proteins have been identified [5], including γ-TURC and the microtubule-stabilizing protein PTRN-1/Patronin/CAMSAP [11,15,40–43]. Additionally, NOCA-1/Ninein is often found to colocalize with the minus ends of microtubules, although it has never been shown to directly bind to minus ends [44]. A previous report found that γ-TURC and NOCA-1 function together in parallel with PTRN-1 to maintain non-centrosomal microtubule arrays in *C. elegans* larval and adult skin [45]. Furthermore, the NOCA-1 h-isoform appears to localize to the membrane ncMTOC in the *C. elegans* adult germline using a palmitoylation site [45]. In the absence of this site, γ-TURC is required to target NOCA-1 to the ncMTOC. We found that both PTRN-1 and NOCA-1 localize to the apical ncMTOC in wild-type embryonic intestinal cells (S4A, B, E Fig). We therefore tested the roles of GIP-1 and MZT-1 in the apical localization of PTRN-1 and NOCA-1. PTRN-1 and NOCA-1(d,e isoforms) appeared to localize normally in GIP-1^gut(-)^ or MZT-1^gut(-)^ embryos (S4C, D, F, G Fig). The NOCA-1 d- and e-isoforms lack the NOCA-1h region containing the characterized palmitoylation site; however, we did not rule out the use of alternative palmitoylation sites, which the d-isoform is predicted to contain (Cys6, [46]).

### AIR-1/Aurora A is required to target TAC-1 and γ-TURC to the centrosome in intestinal cells, but not to the apical ncMTOC

AIR-1/Aurora A is a mitotic kinase that helps activate MTOC function at the centrosome, in part by driving PCM accumulation of targets required for microtubule nucleation and polymerization, such as γ-TURC and TAC-1/TACC [32,47–50]. The phosphorylated, kinase-active form of AIR-1 localizes to the apical ncMTOC along with GIP-1 and TAC-1 [6], suggesting that AIR-1 could similarly regulate the accumulation of these targets at the apical ncMTOC. We first asked whether AIR-1 is required for normal GIP-1 and TAC-1 accumulation at the mitotic centrosome of dividing E8 cells; as predicted from previous studies in other cell types and organisms, average GFP::GIP-1 centrosomal fluorescence was significantly reduced in AIR-1^gut(-)^ cells compared with control cells (532.7 ± 172.8 vs. 1194.6 ± 329.9, p = 1.35 × 10^−7^, Fig 5A-C, see Materials and Methods). We note that centriolar GIP-1 signal appeared to remain upon AIR-1 depletion (Fig 5B). We observed a similar reduction in GFP::TAC-1 levels at the mitotic centrosome of AIR-1^gut(-)^ cells compared to control cells (501.9 ± 91.7 vs. 972.0 ± 463.6, p = 2.88 × 10^−6^, Fig 5D-F), suggesting that AIR-1 can be efficiently depleted prior to ncMTOC formation. In contrast to the E8 mitotic centrosomes, GFP::GIP-1 and GFP::TAC-1 were still recruited to the apical ncMTOC in control and AIR-1^gut(-)^ embryos (Fig 5G-J). These results indicate that AIR-1 is required for GIP-1 and TAC-1 recruitment to intestinal mitotic centrosomes, as is known in other cell types and organisms [48,51], but that AIR-1 is not required for their localization to the apical ncMTOC, further distinguishing the centrosome from the apical ncMTOC.

**Fig. 5.**
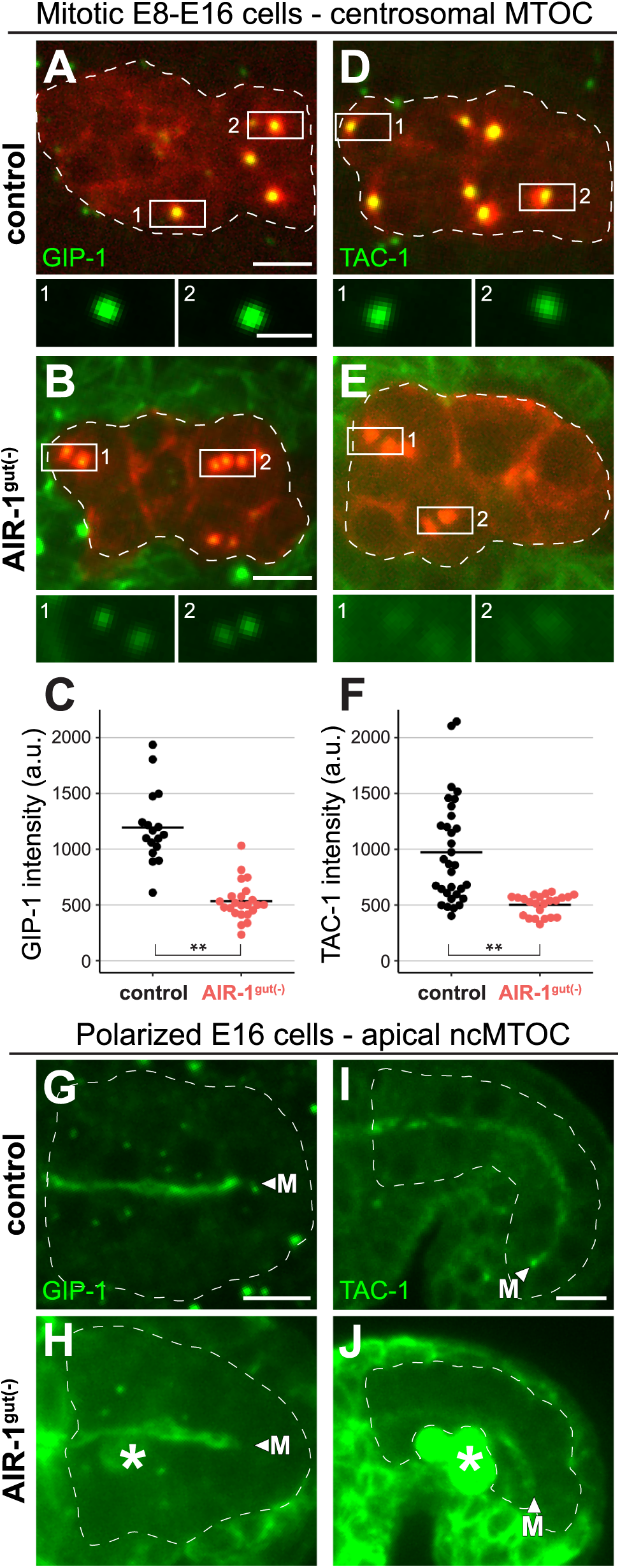
AIR-1 promotes GIP-1 and TAC-1 localization to centrosomes, but not to the apical ncMTOC. Main images are projected optical sections through a portion of the intestinal primordium of *zif*-*1(*-) control (A, D, G, I) and AIR-1^gut(-)^ (B, E, H, J) embryos. Inset images are single optical sections. A-F) Intestine-specific mCherry::TBA-1/α-tubulin marks active centrosomes (enlarged in inset) in dividing E8-E16 cells. Images show endogenous GFP::GIP-1 (A, B) and transgene-expressed GFP::TAC-1 (D, E). Average centrosomal fluorescence intensity is quantified in (C, F; GFP::GIP-1 – control n = 17, AIR-1^gut(-)^ n = 23 centrosomes; GFP::TAC-1 – control n = 32, AIR-1^gut(-)^ n = 23 centrosomes). Note that AIR-1^gut(-)^ images are uncorrected for the high levels of non-degraded AIR-1::ZF::GFP in tissues neighboring the intestinal primordium (see Materials and Methods, S3 Fig), seen here as a green haze and making the significant reduction of GIP-1 and TAC-1 at the centrosome in AIR-1^gut(-)^ cells an underestimate. Asterisks indicate a significant difference from control by two-tailed t-test (p < 1 × 10^−5^). G-J) Images show GFP::GIP-1 (G, H) and GFP::TAC-1 (I, J) in control (G, I) and AIR-1^gut(-)^ (H, J) E16 polarized intestinal primordia. Apical GFP::GIP-1 was observed in 36/36 control and 47/47 AIR-1^gut(-)^ embryos; apical GFP::TAC-1 was observed in 32/32 control and 40/44 AIR-1^gut(-)^ embryos. Apical localization of GFP::TAC-1 is most evident in later comma-stage embryos (I, J). Arrowheads and ‘M’ indicate the apical midline, and asterisks denote AIR-1::ZF::GFP fluorescence in primordial germ cells (H, J). Scale bar is 5 μm in main panels and 2 μm in insets.

### Microtubules remain apically enriched upon depletion of essential microtubule nucleators, anchors, and stabilizers

γ-TuRC and AIR-1 are required to nucleate microtubules at the centrosome in *C. elegans* [20]. We used GIP-1^gut(-)^, MZT-1^gut(-)^, AIR-1^gut(-)^, and [GIP-1; AIR-1^gut(-)^ embryos to test whether these proteins are similarly required to build microtubules at the apical ncMTOC. To monitor microtubules upon depletion, we simultaneously expressed intestine-specific mCherry::TBA-1/α-tubulin along with ‘early’ ZIF-1 (Fig 6A, B) or we expressed ‘late’ ZIF-1 and labeled microtubules ubiquitously with mCherry::TBA-1 from a maternally expressed integrated transgene (Fig 6E). In control embryos, microtubules emanate from the apical surfaces of intestinal cells (Fig 6B, E), an observation that is visualized by plotting mCherry::TBA-1 intensity along a line drawn across the apical midline and quantified by measuring apical enrichment (Fig 6C-G, S3 Fig). Despite severe mitotic defects in GIP-1^gut(-)^ cells revealed by live imaging, microtubules still localize robustly to the apical ncMTOC (Fig 6A, S1 Movie). We see apical microtubule enrichment in MZT-1^gut(-)^, GIP-1^gut(-)^, AIR-1^gut(-)^, and [GIP-1; AIR-1]^gut(-)^ embryos that is not significantly different (Fig 6A-F), and in one case significantly higher (Fig 6G), than in control embryos.

**Fig. 6.**
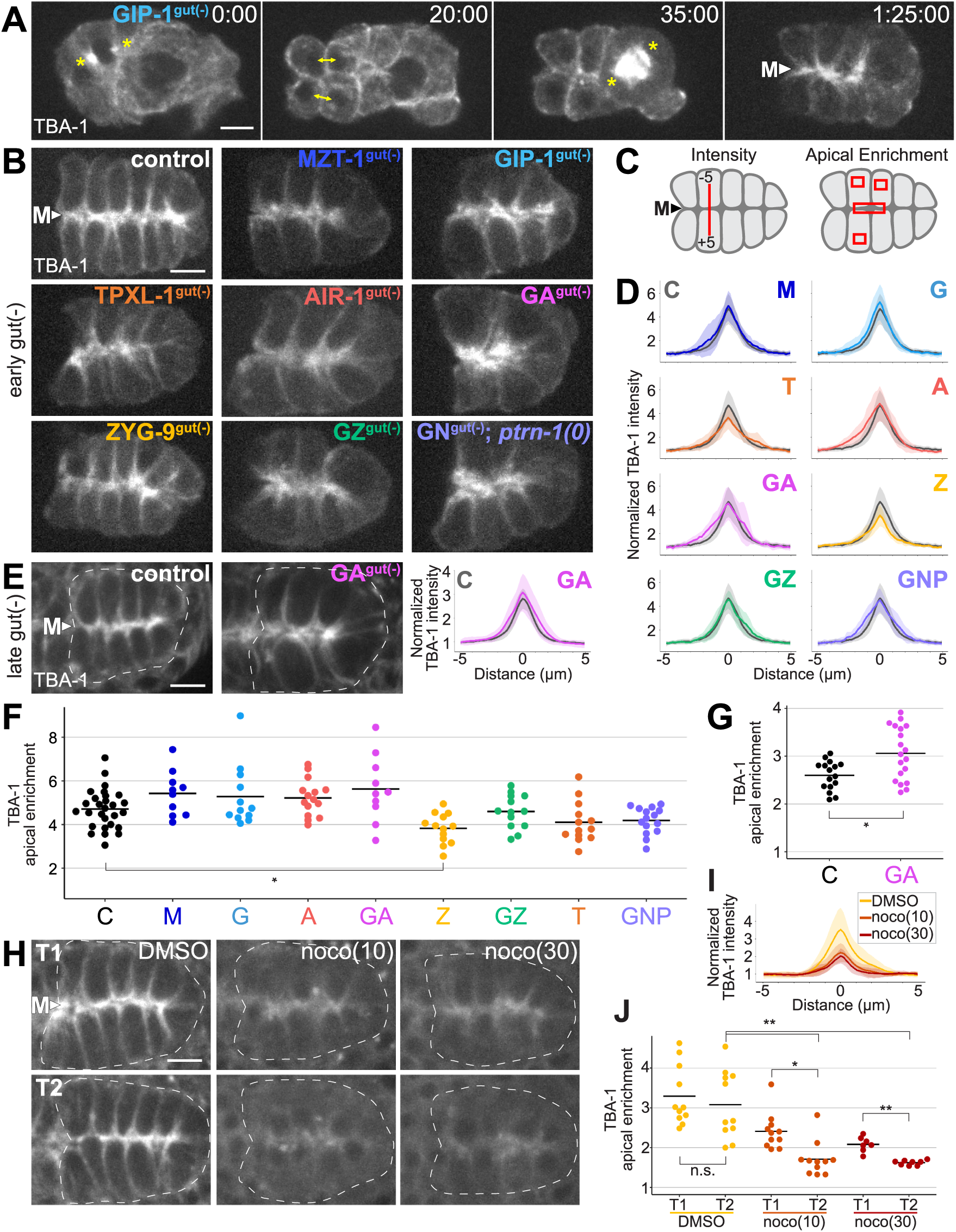
Apical microtubules persist following depletion of GIP-1, MZT-1, and/or AIR-1, and other microtubule regulators A) Still images from S1 Movie. Images show the final mitotic divisions before intestinal polarization and the appearance of the apical ncMTOC in a GIP-1^gut(-)^ embryo. Centrosomes (asterisks) and sister cells (double-headed arrows) are indicated. Note that the posterior of this intestinal primordium contains one rather than 4 cells prior to division, yet apical microtubules still form at the midline (arrowhead and ‘M’). B, E, H) 1.5-μm projected optical sections through the E16 intestinal primordium of embryos with indicated genotypes expressing intestine-specific (B) or maternally expressed (E, H) mCherry::TBA-1/α-tubulin. C) Cartoon summarizing measurements for intensity in D, E, I, and for apical enrichment in F, G, and J. D, E, I) Normalized average TBA-1 pixel intensities along 10-μm line segments drawn across the apical midline of polarized intestinal primordia (n ≥ 10 embryos) for each indicated genotype with early gut(-) (D) and late gut(-) (E), and for control-treated (DMSO) and nocodazole-treated embryos (I). Light shaded regions show standard deviation. Abbreviated genotypes: ‘M’ MZT-1^gut(-)^; ‘G’ GIP-1^gut(-)^; ‘T’ TPXL-1^gut(-)^; ‘A’ AIR-1^gut(-)^; ‘GA’ [GIP-1; AIR-1]^gut(-)^; ‘Z’ ZYG-9^gut(-)^; ‘GZ’ [GIP-1; ZYG-9]^gut(-)^; ‘GNP’ [GIP-1; NOCA-1]^gut(-)^; *ptrn*-*1(0).* ‘C’ *zif*-*1(*-) control profiles are included in gray for reference. F, G, J) Ratio of apical TBA-1 signal as compared to cytoplasmic levels for each indicated genotype with early gut(-) (F), late gut(-) (G), and nocodazole treatment (J). Each dot represents a single embryo of the indicated genotype and black bars indicate mean values for each genotype. No significant differences are observed between genotypes, except for a significant reduction in ZYG-9^gut(-)^ embryos (p = 0.002) and a significant increase in [GIP-1;AIR-1]^late gut(-)^ (p = 0.004) as compared to control. H, I, J) Images are 1.5 μm projected optical sections through the E16 intestinal primordium of embryos treated with DMSO (control), or with 10 μg/mL or 30 μg/mL nocodazole (noco(10) and noco(30)) within 1 minute of treatment (T1) and ten minutes later (T2). mCherry::TBA-1 intensity (I) and apical enrichment (J) is indicated for each case. Each dot represents a single embryo of the indicated genotype and black bars indicate mean values for each genotype. Comparing T1 to T2: DMSO p = 0.54, noco(10) p = 0.002, noco(30) p = 0.0004. Comparing T2 control to T2 treatment: noco(10) p = 0.0003, noco(30) p = 0.0002. In F, G, and J, ^*^p < 0.01, ^**^p < 0.001 by two-tailed t-test. Scale bar is 5 μm for all images.

The surprising finding that known microtubule nucleators are not required to build the majority of microtubules at the apical ncMTOC indicates that other mechanisms or molecular players are required to perform this task. We investigated other known microtubule regulators—the anchoring protein NOCA-1/Ninein, stabilizers PTRN-1/CAMPSAP2 and TPXL-1/TPX2, and the polymerase ZYG-9/chTOG [18,52,53]—to determine if they are required to organize microtubules apically. We found that apical microtubule enrichment was slightly but significantly decreased only upon depletion of ZYG-9, but that even then, microtubules remained apically enriched (Fig 6B, D, F). We were particularly surprised to see grossly normal apical microtubule organization in [GIP-1; NOCA-1]^gut(-)^; *ptrn*-*1(0)* triple mutant embryos, as *noca*-*1* and *ptrn*-*1* are required in parallel to maintain the organization of non-centrosomal microtubule arrays in hypodermal epithelial cells [45]. These results suggest that different MTOCs, and even different ncMTOCs, have distinct molecular and genetic requirements to generate specific microtubule arrays, and that more mechanisms remain to be identified.

### A subset of apical microtubules is perturbed upon depletion of GIP-1

While we found that overall apical organization of microtubules was intact, we explored whether microtubule dynamics were altered upon depletion of microtubule regulators. One possibility is that the majority of microtubules at the apical ncMTOC are stable, persisting from the mitotic divisions prior to polarization. We tested this possibility in two ways. First, we found that upon nocodazole treatment (10 μg/mL and 30 μg/mL), apical microtubule enrichment was significantly reduced both over time and compared to control treated embryos (Fig 6H-J), indicating that apical microtubules can be destabilized. This experiment also demonstrates that our measurement methods (line intensity profiles and apical enrichment) are sensitive enough to detect differences in varying amounts of apical microtubules.

Second, we probed microtubule dynamics at the apical ncMTOC by examining the localization of EBP-2/EB1, a microtubule plus-end-binding protein that associates with growing microtubule plus ends. To do this, we tagged the endogenous *ebp*-*2* locus with GFP using CRISPR/Cas9 to visualize endogenous EBP-2 comets. In control embryos, EBP-2 accumulated at the apical surface, and moved along the apical surface and out along lateral microtubule tracks towards the basal part of the cell, consistent with microtubules growing from the apical ncMTOC (Fig 7A-C, S2 Movie). By contrast, EBP-2 enrichment at the apical surface in GIP-1^gut(-)^ embryos was decreased compared to controls (1.51 vs. 1.67-fold enriched, p = 0.02; Fig 7C). Additionally, we saw a significant decrease in EBP-2 enrichment in [GIP-1; AIR-1]^gut(-)^ compared to AIR-1^gut(-)^ embryos (1.47 vs. 1.64-fold enriched, p = 0.001, Fig 7C), suggesting that the loss of GIP-1 primarily causes the decrease in apical EBP-2 enrichment. Consistent with this observation, we found a significant decrease in the number of EBP-2 comets in [GIP-1; AIR-1]^gut(-)^ compared to AIR-1^gut(-)^ embryos, and in pooled embryos with GIP-1-depleted genotypes compared to pooled embryos with GIP-1(+) genotypes (Fig 7E, Materials and Methods). These results suggest that, unlike at the centrosome, γ-TURC and AIR-1 are not required to build microtubules at the ncMTOC. Instead, γ-TURC is required to build a subset of dynamic microtubules alongside other γ-TuRC-independent microtubules at the apical ncMTOC.

**Fig. 7.**
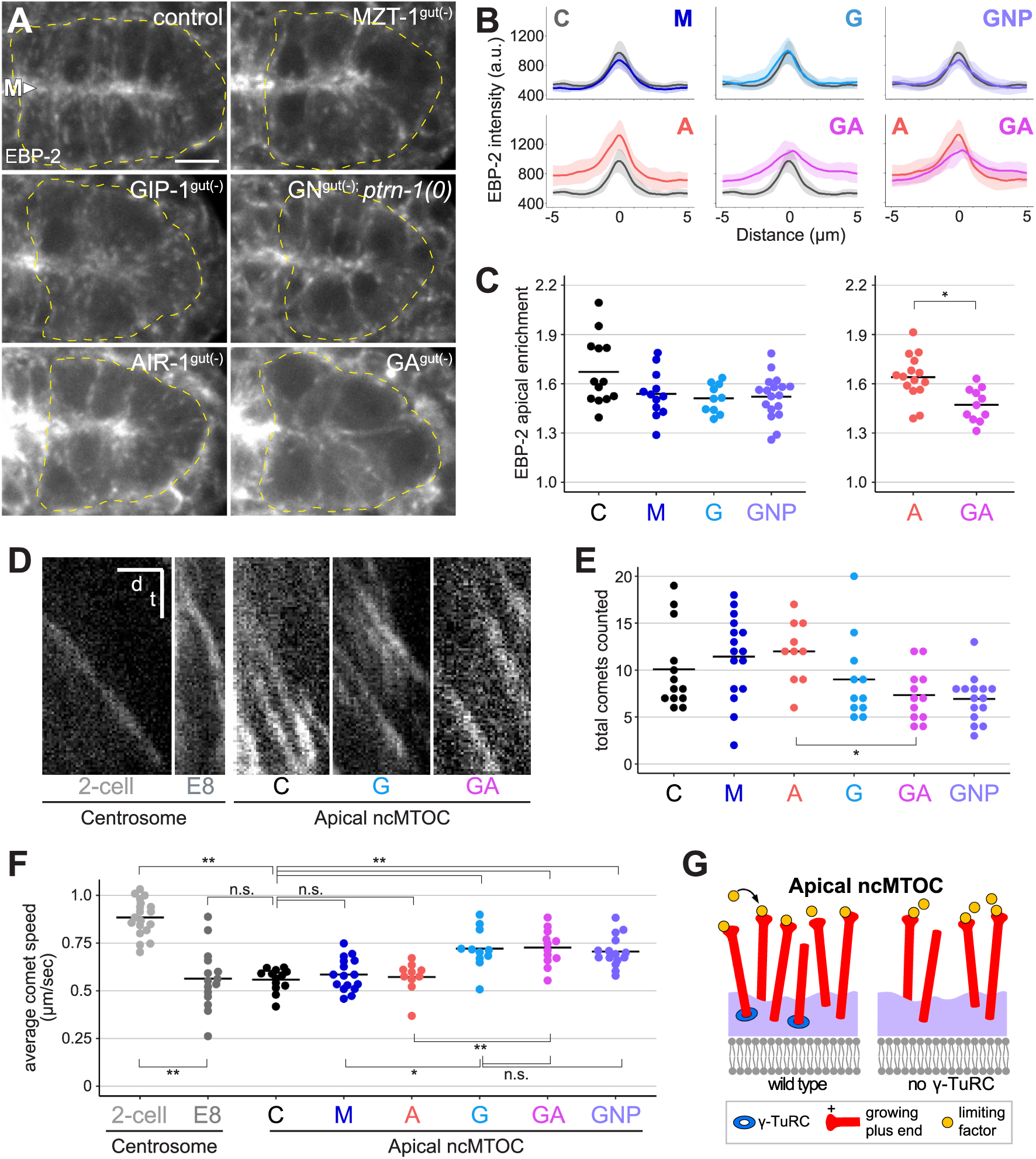
GIP-1 is required for a subset of dynamic microtubules at the apical ncMTOC A) 1.5 μm projected optical sections through the E16 intestinal primordium show EBP-2/EB1 ::GFP localization in live bean-stage embryos of indicated genotypes. Intestinal primordium is outlined by yellow dashed lines; ‘M’ and arrowhead indicate the apical midline. Scale bar is 5 μm. B) Average EBP-2::GFP pixel intensities along line segments drawn through the midline of polarized intestinal primordia (n ≥ 10, see S3 Fig), with abbreviated genotypes indicated (see Fig 6 legend). C) Apical enrichment of EBP-2::GFP signal. Apical EBP-2::GFP is significantly reduced only in GIP-1^gut(-)^ embryos compared to control (p = 0.022, two-tailed t-test), and in [GIP-1;AIR-1]^gut(-)^ compared to AIR-1^gut(-)^ (p = 0.001, twotailed t-test). Apical enrichment of EBP-2::GFP was compared between most genotypes; due to high AIR-1::ZF::GFP background, apical enrichment of EBP-2::GFP in AIR-1^gut(-)^ is only compared to other genotypes that include AIR-1^gut(-)^ (see Materials and Methods). D-F) Rapid time-lapse imaging was used to measure comet speed and number in the intestinal primordia of live embryos of the genotypes indicated. D) Kymographs of EBP-2::GFP comets in indicated structures and stages. Scale bar is 2 sec (t) by 2 μm (d). E) Individual dots show total number of comets crossing two 5 μm lines drawn 3 μm from either side of the apical midline in an individual embryo. F) Average speeds of comets growing from either individual centrosomes (‘2-cell, ‘E8’) or an apical ncMTOC from a single embryo (‘C’, ‘G’, ‘A’, ‘GA’) of the indicated genotype and stage. Total number of comets analyzed per genotype ≥ 30. Comet speed is significantly increased in ‘G,’ ‘GA,’ and ‘GNP’ compared with control embryos (p < 0.0005, two-tailed t-test, see Materials and Methods). In (C-F), ^*^p < 0.01, ^**^p < 0.001 by two-tailed t-test. G) Limiting factor model: Loss of γ-TURC leads to fewer growing microtubules, freeing up a limiting factor that promotes faster microtubule growth at remaining dynamic microtubules.

We next measured the speed of the EBP-2 comets coming from the apical ncMTOC to determine if the dynamics of microtubule growth were altered (Fig 7D, F). In control embryos, apically-derived comets had an average speed of 0.558 μm/sec, which was significantly slower than the reported speeds for comets originating from centrosomes in the early embryo that had been previously labeled with overexpressed EBP-2::GFP [18] or for endogenous early embryo centrosomal comet speeds we measured (‘2-cell’ 0.8884 μm/sec, p = 7.68 × 10^−13^, two-tailed t-test, Fig 7D, F). We also measured comets speeds from centrosomes in the E8-E16 division (‘E8’, 0.563 μm/sec), which had similar speeds to apically-derived comets, and were also significantly slower than comets from centrosomes in the 2-cell embryo (Fig 7D, F), suggesting that cell type, cell size, or centrosome size may influence comet speed [54]. Surprisingly, we found that the speed of apically derived comets was significantly increased relative to controls in GIP-1^gut(-)^ (0.721 μm/sec), [GIP-1; AIR-1]^gut(-)^ (0.726 μm/sec), and [GIP-1; NOCA-1]^gut(-)^; *ptrn*-*1(0)* (0.706 μm/sec) embryos (p <0.0005 for all comparisons, Fig 7D, F). This increase in speed was not observed in AIR-1^gut(-)^ (0.572 μm/sec, p = 0.66) or MZT-1^gut(-)^ embryos (0.585 μm/sec, p = 0.33), and comets in GIP-1^gut(-)^ embryos were significantly faster than in MZT-1^gut(-)^ embryos (p = 0.002).

This lack of an increase in apically derived comet speed following AIR-1 or MZT-1 depletion argues that changes in intestinal morphology caused by too few intestinal cells, which should be nearly identical in AIR-1^gut(-)^, MZT-1^gut(-)^, and GIP-1^gut(-)^ embryos, cannot account for the observed increase in comet speeds in GIP-1^gut(-)^ embryos. These results further demonstrate that MZT-1 is not required for γ-TURC function at the apical ncMTOC. The presence of fewer, faster dynamic microtubules following the depletion of GIP-1 suggests that a limiting microtubule growth factor is normally present at the apical ncMTOC, and that the loss of γ-TuRC-based microtubules frees up that limiting factor, permitting faster growth of γ-TuRC-independent microtubules (Fig 7G).

## Discussion

Using a tissue-specific protein degradation system, we tested the role of factors essential for building microtubules at the centrosome in building microtubules at an ncMTOC. These studies reveal two important findings (Fig 7G, S5 Fig): 1) all MTOCs are not equivalent, with different MTOCs requiring distinct proteins to build and localize microtubules and microtubule regulators, and 2) ncMTOCs can be composed of discrete populations of γ-TuRC-dependent and -independent microtubules.

We demonstrate that our adapted ZIF-1/ZF degradation system is a robust method for depleting endogenous proteins in a specific tissue of interest (the primordial intestinal epithelium), thereby allowing us to probe the function of early essential genes in differentiating tissues. We monitored and characterized the effectiveness and efficiency of endogenous protein depletion by adding both GFP and ZF via CRISPR to genes of interest. Degradation of many of these critical centrosomal proteins during intestinal divisions caused mitotic defects, confirming that targeted proteins were depleted before the apical ncMTOC formed. With this adapted method now validated, future studies can omit the GFP and use ZF-tagged CRISPR alleles, which will permit a broader range of quantitative analyses of GFP markers.

A consequence of early depletion of important centrosomal proteins was fewer intestinal cells, causing architectural defects in the intestinal primordium, such as overall shorter apical midlines. However, the overall reduced midline surface, especially in [GIP-1;AIR-1]^gut(-)^ embryos, does not explain the general upward trend in apical enrichment of α-tubulin we observed; we found no evidence of a correlation between midline length and α-tubulin enrichment (see Materials and Methods). In addition to changes in intestinal geometry, early depletion of centrosomal proteins likely also caused changes in ploidy. While these changes may have impacted zygotic gene expression, they cannot explain, for example, the observed differences among MZT-1^gut(-)^, GIP-1^gut(-)^, and AIR-1^gut(-)^ embryos, which all have similar nuclear numbers and thus likely similar ploidy defects. In sum, differences in microtubule dynamics do not appear to correlate with ploidy or architecture defect severity (Fig 2, 7), indicating that these secondary defects alone cannot account for the changes in microtubule dynamics we observe.

Using the ZIF-1/ZF system, we characterized the predicted *C. elegans* ortholog of MZT1, presenting the first *in vivo* characterization of a MZT1 ortholog in animal development to our knowledge. As in many systems, MZT-1 colocalizes with other γ-TURC components, is required for γ-TURC localization to the mitotic spindle pole, and is essential for viability. Surprisingly, we found that γ-TuRC does not require MZT-1 for its localization to the centrioles and apical MTOC. In addition to localizing γ-TURC, MZT1 is important for nucleation activity of γ-TURC in *Candida* and in human tissue culture cells [38,55]. However, we found that only intestinal GIP-1 depletion, and not MZT-1 depletion, impacted comet number and dynamics, suggesting that the MZT-1 found at the apical MTOC is not important for γ-TURC activity, and may simply be a non-functional component of the γ-TURC complex at these non-PCM sites.

Strikingly, we found that dynamic microtubules were still observed growing from the apical ncMTOC following depletion of GIP-1 and AIR-1, which are essential for centrosomal MTOC activity [20]. This finding indicates that additional mechanisms for generating dynamic microtubules must exist. One exciting possibility is that additional yet undiscovered nucleators exist in the cell. Based on previous studies on centrosomal microtubules, these hypothetical molecules might be unique for building microtubules at ncMTOCs.

Rather than additional nucleators, another possible mechanism could be through the action of microtubule-stabilizing and -anchoring proteins, as has been seen for other ncMTOCs. In fact, the exact role of γ-TURC *in vivo* is not known. The relatively poor nucleation capacity of γ-TURC *in vitro* suggests that factors that activate its nucleation capacity at MTOCs exist *in vivo* [12,56]. Alternatively, the primary function of γ-TURC might not be nucleation, as is suggested by imaging studies of centrosomes from γ-tubulin-depleted *C. elegans* embryos [57]; a large number of microtubules are still found associated with the centrosome following γ-tubulin depletion, but are disorganized relative to the centrioles. These data raise the possibility that γ-TuRC functions in anchoring microtubules onto the PCM, and that perhaps dynamic microtubules at the apical ncMTOC are generated from many different types of stabilized microtubule seeds. For example, proteins like PTRN-1/Patronin/CAMSAP and NOCA-1/Ninein could protect and anchor small microtubule seeds that grow in parallel to the γ-TuRC-based microtubules. Evidence for this type of model has been seen in *Drosophila* oocytes and *C. elegans* skin cells [45,58]. However, our analysis of GIP-1^gut(-)^; NOCA-1^gut(-)^; *ptrn*-*1(0)* triple mutants suggests that in embryonic intestinal cells, the ncMTOC does not require PTRN-1 and NOCA-1/Ninein, even in parallel with GIP-1. Furthermore, microtubules remained organized at the apical MTOC upon depletion of the microtubule polymerase ZYG-9/chTOG and the spindle assembly factor TPXL-1/TPX2, suggesting that additional microtubule regulators remain to be discovered.

A final possibility is that MTOCs could facilitate microtubule growth not by localizing nucleators, but instead by increasing the local tubulin heterodimer concentration. Microtubules can be nucleated *in vitro* in the absence of any additional molecules, depending on the concentration of tubulin. Recent studies of *in vitro* reconstituted PCM suggest that the centrosome might build microtubules in part through the selective concentration of tubulin [53]. ncMTOCs might similarly concentrate tubulin, leading to localized microtubule growth. Two of our findings are consistent with this possibility. First, we observed α-tubulin enrichment at the apical MTOC following microtubule depolymerization with nocodazole, though we cannot distinguish between free tubulin heterodimers and small protected microtubule seeds. Second, we found a small but significant decrease in apical microtubule enrichment in ZYG-9^gut(-)^ embryos, which can concentrate tubulin *in vitro* at SPD-5 condensates [53], and could perhaps play a similar role at the apical ncMTOC.

Our finding that microtubules have increased growth speeds following GIP-1 depletion suggests that a limiting growth factor is normally present at the apical ncMTOC. This limiting factor could be sequestered by γ-TURC itself, or it could be limiting because of competition for it among the large number of growing microtubules. In the first case, we would expect loss of γ-TURC to release this factor and cause both increased microtubule growth speed and comet number. However, we observed fewer, faster comets and decreased apical EBP-2 enrichment upon GIP-1 depletion. These observations are more consistent with a model in which loss of γ-TURC leads to fewer growing microtubules, thereby increasing the availability of a limiting factor to growing microtubules and allowing faster growth (Fig 7G). This limiting factor could be tubulin heterodimers themselves, however, the mechanisms for concentrating a pool of tubulin at a membrane are completely unknown.

Finally, we found that γ-TURC and AIR-1 are not required to form the majority of apical microtubules, raising the question of why these proteins so specifically localize there as a new ncMTOC is being established. One possibility is that the main function of localizing γ-TURC and AIR-1 to the apical ncMTOC is to effectively remove them from the centrosome at the end of mitosis as centrosomal MTOC function is attenuated. We hypothesize that the ability to remove microtubules from the centrosome is an important step in mitotic exit, as hyperactive MTOC function at the centrosome has been linked to cancer [59–61]. Creating a sink for centrosomal microtubule regulators at an alternative site in the cell would provide a quick and effective way of maintaining the inactivation of MTOC function at the centrosome. Different cell types require a large variety of specific patterns of microtubule organization, and future work will be critical to discover the additional molecular players and mechanisms that contribute to the formation and function of different types of MTOCs.

## Materials and Methods

All data used for quantitative analyses are included as S1 Data. Image files are available upon request.

### C. elegans strains and maintenance

Nematodes were cultured and manipulated as previously described [62]. Experiments were performed using one- or two-day old adults. The strains used in this study are as follows:

| Strain Name | Genotype | Fig | Source |
| --- | --- | --- | --- |
| JLF22 | *sec-5(xn51[sec-5::zf::yfp+LoxP::unc-119::loxP])* II; *zif-1(gk117)* III | 1F | This study |
| JLF23 | *zif-1(gk117)* III; *axIs1462[pie-1p::gfp::pie-1::pie-1 3'UTR + unc-119(+)]* | 1D | This study |
| JLF36 | *zuls278[pie-1p::cherry::tba-1::pie-1 3'UTR]; zif-1(gk117)* III | 6H | This study |
| JLF38 | *gip-1(wow5[zf::gfp::gip-1]) zif-1(gk117) III; wowEx10* | 1H-J, 2G, 6A,B, S1B, S1 Movie | This study |
| JLF43 | *zif-1(gk117) III; wowEx10* | 6B | This study |
| JLF48 | *gip-1(wow5[zf::gfp::gip-1]) zif-1(gk117) III; wowEx15[ges-1p::zif-1]* | S2I | This study |
| JLF109 | *zyg-9(wow13[zf::gfp::zyg-9]) II; zif-1(gk117) III; wowEx10* | 6B | This study |
| JLF112 | *zif-1(gk117) III; wowEx28* | 3I | This study |
| JLF113 | *gip-1(wow5[zf::gfp::gip-1]) zif-1(gk117) III; wowEx28* | 3J | This study |
| JLF126 | *gip-1(wow5[zf::gfp::gip-1]) zif-1(gk117) III; wowEx34* | S1B | This study |
| JLF155 | *zif-1(gk117) III, outcrossed 6x* | text | This study |
| JLF174 | *zif-1(gk117) stls10220 [end-1p::histone1::mcherry] III; air-1(wow14[air-1::zf::gfp]) V; wowEx34* | 2B | This study |
| JLF175 | *zif-1(gk117)/eT1 III; air-1(wow14[air-1::zf::gfp])/eT1[nls267[myo-2p::gfp]] V; wowEx10* | 1O,P, 6B, S1C | This study |
| JLF189 | *zif-1(gk117) III; air-1(wow14[air-1::zf::gfp]) V; wowEx34* | S1C | This study |
| JLF193 | *zif-1(gk117) stls10220/eT1 III; air-1(wow14[air-1::zf::gfp])/eT1[nls267] V; wowEx10* | 2B | This study |
| JLF210 | *zuls278; gip-1(wow5[zf::gfp::gip-1]) zif-1(gk117) III; air-1(wow14[air-1::zf::gfp]) V; wow/s3[spontaneous wowEx34 integrant]/+* | 6E | This study |
| JLF226 | *gip-1(wow3[gfp::gip-1]) zif-1(gk117) III;* | 3F, 4A-C, | This study |
|  | *wowEx10* | 5A,F |  |
| JLF227 | *gip-1(wow5[zf::gfp::gip-1]) zif-1(gk117) stls10220 III; wowEx34* | 2B | This study |
| JLF229 | *ItSi540[noca-1p::noca-1(de)::superfolder-gfp] II; zif-1(gk117) III; wowEx10* | S4E | This study/KO |
| JLF230 | *bcls104[pie-1p::gfp::tac-1]; zif-1(gk117) III; wowEx10* | 5C,H | This study |
| JLF231 | *bcls104; zif-1(gk117)/eT1 III; air-1(wow14[air-1::zf::gfp])/eT1[nls267] V; wowEx10* | 5D,I | This study |
| JLF232 | *gip-1(wow5[zf::gfp::gip-1]) zif-1(gk117) III; air-1(wow14[air-1::zf::gfp]) V; wowls3/+* | S1D, S2 | This study |
| JLF233 | *gip-1(wow5[zf::gfp::gip-1]) zif-1(gk117) stls10220 III; wowEx34* | 2B | This study |
| JLF234 | *gip-1(wow5[zf::gfp::gip-1]) zif-1(gk117) stls10220 III; air-1(wow14[air-1::zf::gfp]) V; wowEx34* | 2A,B | This study |
| JLF235 | *zif-1(gk117) III; ptrn-1(wow4[ptrn-1::gfp]) X; wowEx10* | S4B | This study |
| JLF237 | *gip-1(wow3[gfp::gip-1]) zif-1(gk117) III; air-1(wow14[air-1::zf::gfp])/oxTi878[vha-6p::gfp] V; wowEx10* | 5B,G | This study/CGC |
| JLF239 | *mzt-1(wow23[zf::gfp::mzt-1]) I; gip-1(wow3[gfp::gip-1]) zif-1(gk117) III; wowEx10* | 3H, 4D-F | This study |
| JLF245 | *gip-1(wow5[zf::gfp::gip-1]) zif-1(gk117) III; air-1(wow14[air-1::zf::gfp])/oxTi878 V; wowEx10* | 6B, S1D | This study |
| JLF247 | *zif-1(gk117) stls10220 III; air-1(ok571)/oxTi878 V; wowEx10* | 2B | This study/CGC |
| JLF249 | *gip-1(wow5[zf::gfp::gip-1]) zif-1(gk117) stls10220 III; wowEx10* | 2B | This study |
| JLF255 | *mzt-1(wow23[zf::gfp::mzt-1]) I; ItSi540 II; zif-* | S4G | This |
|  | *1(gk117) III; wowEx10* |  | study/KO |
| JLF257 | *ItSi540 II; gip-1(wow5[zf::gfp::gip-1]) zif-1(gk117) III; wowEx10* | S4F | This study/KO |
| JLF259 | *mzt-1(wow23[zf::gfp::mzt-1]) I; zif-1(gk117) III; ptrn-1(wow4[ptrn-1::gfp]) X; wowEx10* | S4D | This study |
| JLF260 | *mzt-1(wow23[zf::gfp::mzt-1]) I; stls10220 zif-1(gk117) III; wowEx10* | 2B | This study/CGC |
| JLF268 | *gip-1(wow5[zf::gfp::gip-1]) zif-1(gk117) stls10220 III; air-1(wow14[air-1::zf::gfp])/oxTi878 V; wowEx10* | 2A,B | This study/CGC |
| JLF273 | *ebp-2(wow47[ebp-2::gfp]) II; zif-1(gk117) III; wowEx10* | 7A,D, S2<br>Movie | This study |
| JLF274 | *ebp-2(wow47[ebp-2::gfp]) II; zif-1(gk117)/eT1 III; air-1(wow14[air-1::zf::gfp])/eT1[nls267] V; wowEx10* | 7A, S2<br>Movie | This study |
| JLF275 | *ebp-2(wow47[ebp-2::gfp]) II; gip-1(wow5[zf::gfp::gip-1]) zif-1(gk117) III; wowEx10* | 7A,D, S2<br>Movie | This study/CGC |
| JLF298 | *ebp-2(wow47[ebp-2::gfp]) II; gip-1(wow5[zf::gfp::gip-1]) zif-1(gk117) III; air-1(wow14[air-1::zf::gfp]) V* | 7A,D, S2<br>Movie | This study |
| JLF304 | *ebp-2(wow47[ebp-2::gfp]) II; zif-1(gk117)/eT1[nls267] III; wowEx10* | 7A,D, S2<br>Movie | This study |
| JLF318 | *mzt-1(wow23[zf::gfp::mzt-1]) I; zif-1(gk117) III; wowEx10* | 1L,M, 3E,<br>6B, S1E | This study |
| JLF343 | *gip-1(wow5[zf::gfp::gip-1]) zif-1(gk117) III; ptrn-1(wow4[ptrn-1::gfp]) X; wowEx10* | S4C | This study |
| JLF357 | *mzt-1(wow23[zf::gfp::mzt-1])/hT2[qls48] I; zif-1(+)/hT2 III* | text | This study |
| JLF358 | *zyg-9(wow13[zf::gfp::zyg-9]) II; gip-* | 6B | This study |
|  | *1(wow5[zf::gfp::gip-1]) zif-1(gk117) III; wowEx10* |  |  |
| JLF366 | *zif-1(gk117) III; ddls6[pie-1p::gfp::tbg-1]; wowEx10* | 3L | This study/CGC |
| JLF367 | *gip-1(wow5[zf::gfp::gip-1]) zif-1(gk117)/ qC1 [qls26] III; noca-1(wow11[noca-1::zf::gfp])/oxTi878 V; ptrn-1(lt1) X; wowEx10* | 6B | This study/CGC |
| JLF373 | *mzt-1(wow51[gfp::mzt-1])/hT2[qls48] I; zif-1(+)/hT2 III* | text | This study |
| JLF374 | *ebp-2(wow47[ebp-2::gfp]) II; gip-1(wow5[zf::gfp::gip-1]) zif-1(gk117)/ qC1 [qls26] III; noca-1(wow11[noca-1::zf::gfp])/oxTi878 V; ptrn-1(lt1) X; wowEx10* | 7A, S2 Movie | This study/CGC /KO |
| JLF375 | *mzt-1(wow51[gfp::mzt-1]) I; zif-1(gk117) III; wowEx10* | 3C | This study |
| JLF380 | *mzt-1(wow23[zf::gfp::mzt-1]) I; zif-1(gk117) III; ddls6; wowEx10* | 3N | This study/CGC |
| JLF381 | *gip-1(wow5[zf::gfp::gip-1]) zif-1(gk117) III; ddls6; wowEx10* | 3M | This study/CGC |
| JLF382 | *tpxl-1(wow34[zf::gfp::tpxl-1]) I; zif-1(gk117) III; wowEx10* | 6B | This study |
| JLF383 | *mzt-1(wow51[gfp::mzt-1]) I; gip-1(wow5[zf::gfp::gip-1]) zif-1(gk117) III; wowEx10* | 3D | This study |
| JLF385 | *mzt-1(wow23[zf::gfp::mzt-1]) I; gip-1(wow25[tagRFP::gip-1]) zif-1(gk117) III* | 3B | This study |
|  | *mzt-1(wow23[zf::gfp::mzt-1]) I; zif-1(gk117) III; wowEx28* | 3K | This study |
| JH2015 | *axls1462; unc-119(ed3) III* | 1C | CGC |
| FT1523 | *sec-5(xn51[sec-5::zf::yfp + LoxP::unc-* | 1E | CGC/JN |
|  | *119::loxP*) II; *unc-119(ed3)* III |  |  |
| N2 | Wild type | text | CGC |

### CRISPR

Two CRISPR editing methods were used to generate knock-in insertions of the ZF::GFP or GFP tag. *gip*-*1(wow3[gfp::gip*-*1]*, *gip*-*1(wow5[zf::gfp::gip*-*1]*) and *ptrn*-*1(wow4[ptrn*-*1::gfp])* were constructed using repair templates with short homology arms (SHA) and PCR-based screening to test for insertions into the endogenous loci [63]. All other CRISPR-based insertions were achieved using a modified self-excising cassette (SEC) into which the ZF-tag had been inserted N-terminal to GFP (plasmid JF250) [64]. The SEC-derived CRISPR alleles thus also contain additional tags: 3xFLAG for GFP alleles and 3xMyc for TagRFP-T alleles. Cas9 and sgRNAs were delivered using plasmid pDD162 into which the appropriate sgRNA sequence had been added with the Q5 Site-Directed Mutagenesis Kit (NEB). The Cas9/sgRNA plasmid, repair template, and appropriate selection markers were injected into N2 or *zif*-*1(gk117)* mutant one-day-old adult worms. Worms were recovered and processed according to published protocols [63,64]. Successfully edited worms were outcrossed at least 2 times before being used for subsequent experiments. sgRNA and homology arm sequences are listed in S1 Table.

### Embryonic viability

To assess embryonic viability, 20-30 young adult hermaphrodites of each genotype were singled onto small plates and allowed to lay for 4 hours at 20°C, and then adults were removed and eggs were counted. After three days at 20°C, the number of surviving worms present on each plate was counted, and viability for each plate was calculated as the total number of L4s and adults divided by the number of eggs. When comparing N2 and *zif*-*1(*-) (JLF155) lethality, two N2 plates had more surviving worms at day three than the number of eggs initially counted, and those plates were omitted.

### ZF/ZIF-1 degradation

The ZF/ZIF-1 system was executed as previously described with the following modifications. ZF::GFP tags were inserted into endogenous loci in a *zif*-*1(gk117)* mutant background using CRISPR editing techniques (see above). Exogenous ZIF-1 was expressed from either extrachromosomal or integrated arrays (see below), which were generated by injecting 50 ng/μL of each plasmid and of a co-injection marker (either pRF4[*rol*-*6(d)*] at 100 ng/μL or *myo-2*p::*mcherry* at 2.5 ng/μL) into *zif*-*1(gk117)* mutant animals, with pBS to inject at a minimum concentration of 150 ng/μL. For comparisons across multiple genotypes, the same array was introduced by mating.

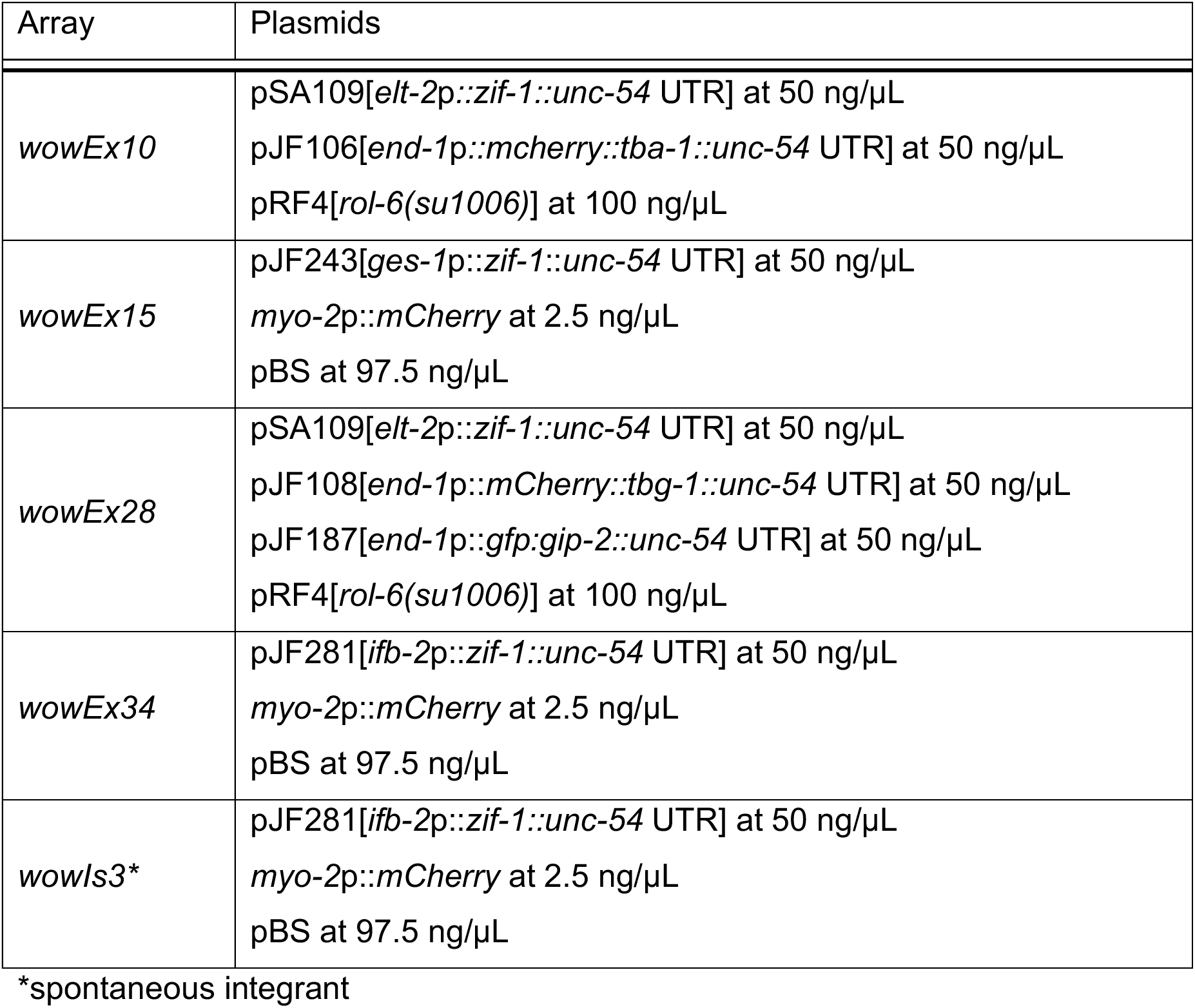

To test the efficiency of AIR-1 depletion—specifically, to test whether zygotic expression of AIR-1::ZF::GFP provided AIR-1 activity in AIR-1^gut(-)^ embryos—we generated *air*-*1(0)* embryos with only maternally provided AIR-1::ZF::GFP, which was degraded in the intestinal primordium by ZIF-1 expression. Our genetic strategy was as follows: We used the *air*-*1(ok571)* allele, a deletion that removes a large portion of the AIR-1 kinase domain and is a putative null allele. We crossed *zif*-*1(gk117) stIs10220[end*-*1*p::*his1::mcherry]; air*-*1(ok571)/oxTi878[vha*-*6*p::*gfp]; wowEx10[elt*-*2*p::*zif*-*1; end*-*1* p::*mcherry::tba*-*1*] hermaphrodites with *zif*-*1(gk117) stIs10220; air*-*1(wow14[air*-*1::zf::gfp]*) males to generate F1 “*air*-*1::zf::gfp/*(0)” hermaphrodites, and scored their F2 embryos. We had no markers to distinguish embryonic genotypes, so we scored all progeny and assumed Mendelian segregation of genotypes: 25% *air*-*1::zf::gfp* homozygotes, 50% *air*-*1::zf::gfp/*(0) heterozygotes, and 25% *air*-*1(0)* homozygotes, all with maternally expressed AIR-1::ZF::GFP. Consistent with this predicted distribution, we saw many larvae with perturbed gonad development, a hallmark of the *air*-*1(0*) phenotype [65]. We denote these pooled embryos as AIR-1^*^ in Fig 2. If zygotic expression of AIR-1 was providing activity, we would expect to observe a change in the distribution of intestinal nuclear number in AIR-1^gut(-)^ compared with AIR-1^gut(-)*^. However, intestinal nuclear number in AIR-1^gut(-)*^ and AIR-1^gut(-)^ embryos were indistinguishable, suggesting that zygotic expression of AIR-1::ZF::GFP provides little or no AIR-1 rescuing activity, and that AIR-1::ZF::GFP depletion is highly efficient.

We used α-GFP immunostaining to test how completely ZF::GFP-tagged proteins are degraded in the intestinal primordium (see S2 Fig). We stained [GIP-1; AIR-1]^gut(-)^ embryos (JLF232) in which heterozygous *ifb-2*p*::zif*-*1(wowIs3)* causes intestine-specific degradation of both proteins in embryos that inherit the transgene. GFP fluorescence survives the staining protocol, so we asked how the perduring GFP signal correlates with Cy3 signal from α-GFP staining. Most JLF232 embryos fell into two categories: strong Cy3 and GFP signal at centrosomes or the apical membrane (n>40), or no localized Cy3 nor GFP signal in the intestinal primordium (n = 30). As α-GFP antibody staining is highly sensitive, this result suggests that GIP-1 and AIR-1 are both degraded quickly (prior to GFP maturation) and efficiently, and that the loss of visible GFP in live embryos accurately indicates efficient ZF-tagged protein degradation.

### Generating EBP-2::GFP; [GIP-1;AIR-1]^gut(-)^ embryos

We did not find a balancer that allowed us to maintain an EBP-2::GFP; [GIP-1;AIR-1]^gut(-)^ strain. We thus used the following cross strategy to generate these embryos: *ebp*-*2(wow47*[*ebp*-*2::gfp])*; *zif*-*1(gk117)/eT1[nIs267]*; *wowEx10[elt*-*2*p::*zif*-*1]* hermaphrodites were crossed with JLF298 males to generate *ebp*-*2::gfp*; *zf::gfp::gip*-*1 zif*-*1(gk117)/eT1[nIs267]*; *air*-*1::zf::gfp/eT1*; *wowEx10* F1 hermaphrodites. F1 cross progeny were identified by PCR genotyping for *air*-*1::zf::gfp* (*air*-*1* F: GAACGTCTCCCACTTGTTGACATC, ZF R: TTTTTCTACCGGTACCCTCGG). F2 progeny with the *wowEx10* array and without the *eT1* balancer were selected (*ebp*-*2::gfp*; *zf::gfp::gip*-*1 zif*-*1(gk117); air*-*1::zf::gfp; wowEx10). zf::gfp::gip*-*1* was frequently heterozygous, likely due to recombination between *eT1* and *zf::gfp::gip*-*1* in the F1 mother. Thus, to ensure we were looking at EBP-2::GFP; [GIP-1;AIR-1]^gut(-)^ embryos, we only scored embryos with four intestinal nuclei (see Fig 2).

### Nocodazole Treatments

JLF36 embryos were treated with nocodazole as has been previously described [6]. Briefly, trypan blue-coated embryos were affixed to poly-lysine-coated coverslips and submerged in embryonic growth medium (EGM) [66] containing either 0.1% DMSO or 10 or 30 μg/mL nocodazole. E16-stage embryos with their dorsal surfaces facing the coverslip were selected for analysis. The eggshell and vitelline membrane were punctured using a Micropoint nitrogen dye laser (Andor), allowing the EGM + DMSO or nocodazole to reach the embryo. Embryos were imaged immediately following puncturing (T1, 10-45 seconds later) and then again after 10 minutes (T2).

### Immunofluorescence

Embryos were fixed and stained as previously described [23]. Briefly, embryos of the appropriate stage were collected and adhered to a poly-lysine-coated slide with a Teflon spacer and covered with a coverslip. Embryos were fixed by freeze-crack followed by 100% MeOH for 5 minutes. Embryos were rehydrated in PBS and incubated with rabbit α-GFP primary antibody (Abcam, 1/200) or mouse α-PAR-3 [28] either overnight at 4°C or for 1 hour at 37°C. Embryos were washed in PBT and incubated with the appropriate CY3-conjugated secondary antibodies (Jackson Immunoresearch Laboratories, 1/200) and 100 ng/mL DAPI (Sigma) overnight at 4°C or for 1 hour at 37°C and mounted under coverslips in Vectashield (Vector Laboratories). At least 20 mutant embryos were scored for analysis of mutant phenotypes.

### Microscopy

Embryos for microscopy were dissected from gravid hermaphrodites incubated for 4-4.5 hours in M9 (at 20°C for EBP-2::GFP comet speed, at 25°C for EBP-2::GFP enrichment and at room temperature for all other experiments). For live imaging, samples were mounted on a pad made of 3% agarose dissolved in M9. Live imaging was performed on a Nikon Ti-E inverted microscope (Nikon Instruments) using a 60x Oil Plan Apochromat (NA = 1.4) or 100x Oil Plan Apochromat (NA = 1.45) objective and controlled by NIS Elements software (Nikon). Images were acquired with an Andor Ixon Ultra back thinned EM-CCD camera using 488 nm or 561 nm imaging lasers and a Yokogawa X1 confocal spinning disk head equipped with a 1.5x magnifying lens. For live imaging, images were taken at a sampling rate of 0.5 μm. Images were processed in NIS Elements, the Fiji distribution of ImageJ (‘Fiji’) [67,68], or Adobe Photoshop. Fixed images were obtained using a 60x Oil Plan Apochromat objective (NA =.1.4) on either the above system or a Nikon Ni-E compound microscope with an Andor Zyla sCMOS camera.

### Image Quantification

#### 1. Analysis Considerations

We observed that different ZF::GFP-tagged alleles produced different levels of background signal in the intestinal primordium following ZIF-1-mediated depletion. This background signal was likely due to out-of-focus light produced by the non-degraded ZF::GFP-tagged proteins in neighboring germ cell precursors, muscle, and pharyngeal cells. In support of this reasoning, we observed significantly different amounts of out-of-focus light in a region directly adjacent to, but outside of embryos of different genotypes. Average external signal in the 488nm channel (400 ms exposure, 50% laser power): *zif*-*1* control: 10.2, GIP-1^gut(-)^: 34.3, MZT-1^gut(-)^: 53.9, AIR-1^gut(-)^: 246.9, [GIP-1;AIR-1]^gut(-)^: 309.8, [GIP-1;NOCA-1]^gut(-)^; *ptrn*-*1(0)*: 58.0. All of these are significantly different from the control (p < 0.00005, two-tailed t-test); this variability in background precluded most direct comparisons of GFP marker intensity in these different genotypes, except for EBP-2::GFP. When we made the same measurements when EBP-2::GFP was present (300 ms exposure, 25% laser power), we observed the following: *zif*-*1* control: 137.1, GIP-1^gut(-)^: 152.6, MZT-1^gut(-)^: 142.7, AIR-1^gut(-)^: 210.5, [GIP-1;AIR-1]^gut(-)^: 222.7, [GIP-1;NOCA-1]^gut(-)^; *ptrn*-*1(0)*: 133.1. By pairwise t-test comparisons, the two AIR-1 genotypes [GIP-1;AIR-1]^gut(-)^ and AIR-1^gut(-)^ are significantly different from each of the other four genotypes but not from each other. None of the other four genotypes are significantly different from each other, allowing us to break the genotypes into two groups for comparison: genotypes with AIR-1^gut(-)^ and genotypes with no tagged AIR-1 (Fig 7B, C). For genotypes that did not meet this quantitative cutoff, our comparisons were qualitative. These differences in background did not impede our analysis of EBP-2::GFP comet number and speed, since comet signal was visible above background signal in all genotypes analyzed. Additionally, because all mCherry signal was coming only from TBA-1 transgenes, there were no differences in background mCherry signal between genotypes, allowing for unrestricted comparisons.

Embryos of the appropriate stage with their dorsal side oriented toward the coverslip were chosen for analysis. For all images, three rectangles of variable size were drawn to measure average slide background signal. An R script was used to calculate background-subtracted pixel values and average pixel intensity for each ROI.

#### 2. Quantification of GFP depletion

Pre-bean- and bean-stage embryos were used for analysis. Using Fiji, *end-1*p-driven mCherry::TBA-1 signal was used to select the brightest plane to define the intestinal midline. A sum Z-projection of 3 slices around this plane was used for the analysis. GFP depletion in the intestinal primordium in the different ZF::GFP strains was quantified in two different ways. First, the average fluorescence was measured in a 10-μm-wide box of variable length drawn inside the intestinal primordium (gut (G), S3A Fig). We accounted for out-of-focus light from ZF::GFP in neighboring cells by subtracting the average pixel intensity inside a 2-μm-wide box of variable length drawn outside the embryo (external (X), S3A Fig). Intestinal autofluorescence (‘auto,’ S3A Fig) was estimated by averaging the green signal intensity in the gut region of control *zif*-*1* embryos that lack GFP. Perduring ZF::GFP signal in the intestinal primordium was calculated for GIP-1^gut(-)^, MZT-1 ^gut(-)^, and AIR-1 ^gut(-)^ (G_ZF:ex10_, S3A Fig) and compared to the respective non-degraded GIP-1^gut(+)^, MZT-1 ^gut(+)^, and AIR-1 ^gut(+)^ siblings lacking the array (G_ZF_, S3A Fig). Percent GFP depletion was calculated as indicated in S3A Fig, left.

Second, a 1-μm-wide line segment was drawn across the intestinal midline and pixels within 5 μm of the midline were selected to generate a line profile (S3B Fig). For each genotype, a mean profile was calculated by averaging the intensity value for each distance from the midline. The cytoplasmic region (‘cyto,’ S3B Fig) of each profile was defined as the points between 2.5 and 5 μm from the midline. The peak region was defined for each protein as the distance from the midline at which GFP intensity is halfway between the mean cytoplasmic value and the maximum value in the averaged profile. We compared the mean peak and cytoplasmic intensity values to assess apical enrichment for each genotype (paired Student’s t-test, S3D Fig). In addition, we compared mean peak or mean cytoplasmic intensity values between paired gut(-) and gut(+) genotypes (Welch two sample t-test, S3D Fig).

#### 3. Quantification of centrosomal protein accumulation in AIR-1^gut(-)^ mutants

Despite the high background in AIR-1^gut(-)^ embryos (see above), we were able to measure and detect a clear difference in centrosomal TAC-1 and GIP-1 even with no correction for genetic background (AIR-1::ZF::GFP signal from outside the intestinal primordium). Embryos in the E8-E16 division were chosen for analysis. Centrosomes with strong mCherry::TBA-1 localization were considered active MTOCs and their average GFP intensity was measured. A 7-pixel diameter circular ROI was placed manually to measure the average intensity of GFP::GIP-1 and GFP::TAC-1 at the Z-section of maximum GFP intensity at active centrosomes in *air-1(+)* and AIR-1^gut(-)^ backgrounds. Signal intensities were compared using Welch two-sample t-tests.

#### 4. Quantification of apical α-tubulin and EBP-2 enrichment

Pre-bean- and bean-stage embryos were chosen for analysis. Using Fiji, *end-1*p-driven mCherry::TBA-1 signal was used to select the brightest plane to define the intestinal midline. A sum Z-projection of 3 slices around this plane was used for the remaining analysis. 2-μm-wide boxes in apical and cytoplasmic regions were drawn by hand to define ROIs (S3A Fig, right). Apical enrichment was then calculated by dividing the mean apical intensity value by the mean cytoplasmic intensity value for each image (S3A Fig). Welch two-sample t-tests were used to compare enrichment values between genotypes. A 1-μm-wide line segment was drawn and pixels within 5 μm of the midline were selected to generate a line profile (S3B Fig). Profile plots and plots of apical enrichment were generated in R with ggplot2 [69] and ggbeeswarm (https://cran.r-proiect.org/web/packages/ggbeeswarm), respectively; because mCherry::TBA-1 was expressed from an extrachromosomal array, each profile was normalized by dividing each pixel intensity value by the average intensity in the region 2.5 μm to 5 μm from the midline to correct for different levels of array expression. EBP-2::GFP was a CRISPR allele, so profile plots were not normalized. For each genotype, the mean profile was calculated by averaging the intensity value for each distance from the midline.

We tested whether shorter midline length (and thus a smaller surface area for the ncMTOC) might correlate with higher microtubule enrichment. Indeed, the average midline length is shorter in [GIP-1;AIR-1]^late gut(-)^ embryos than in controls (11.0 μm versus 14.5 μm, t-test p = 3.6×10^−6^), and average apical microtubule enrichment is higher in [GIP-1;AIR-1]^late gut(-)^ than in controls (3.1 versus 2.6, t-test p = 0.004). However, apical enrichment does not correlate with midline length for control and [GIP-1;AIR-1]^late gut(-)^ embryos (linear fit; control R^2^ = 0.0015 and p = 0.90, [GIP-1;AIR-1]^late gut(-)^ R^2^ = 1.9×10^−5^ and p = 0.99), suggesting that changes in midline length do not affect our microtubule enrichment results.

#### 5. Quantification of EBP-2 comets

Time-lapse images of pre-bean and bean-stage embryos were obtained using a 100X objective (see Microscopy). All time-lapse images were processed to correct for photobleaching prior to analysis using the Bleach Correction plugin for Fiji. *end*-*1*p-driven mCherry::TBA-1 signal was used to identify the intestinal midline to guide image acquisition. To count EBP-2 comets coming from the midline, a 5-μm line was overlaid 3 μm from the midline on either side (S3C Fig, top). The number of comets that crossed each line within 10 seconds was counted manually. To measure comet speed, kymographs were generating using the Multi Kymograph plugin of Fiji with a 5-pixel line width (S3C Fig, bottom). Within each kymograph, the slope of one comet was measured (S3C Fig). For E16 embryos with only one measured comet, images were re-analyzed to include at least two measured comets, so that each embryo’s average comet speed was based on two or more measured comets. 2-cell embryos with only one measured comet were excluded from analysis. Comet speed was calculated using R. Comet count and speed were compared between genotypes using Welch two-sample t-tests. In comparing comet number between genotypes, we did not observe a significant decrease in GIP-1^gut(-)^ compared to control alone, but when we pooled embryos from the three GIP-1^gut(-)^ genotypes (G, GA, GNP) and compared them to pooled embryos with wild-type *gip*-*1* (C, A, M), we saw a significant difference between the groups: pooled(GIP-1^gut(-)^) had an average of 7.7 ± 3.3 comets per embryo, and pooled(*gip*-*1(+)*) had an average of 11.1 ± 4.1 comets per embryo (p = 0.0001, two-tailed t-test). Importantly, none of the individual genotypes within a pooled group was significantly different from the others by paired t-tests. All of the gut(-) genotypes with EBP-2::GFP measured in Fig. 7 have excess α-tubulin added from the *zif*-*1* -expressing transgene *wowEx10.* To test if overexpressed α-tubulin was influencing our comet speed results, we measured comet speed in control and GIP-1^late gut(-)^ embryos which do not overexpress tubulin. In the absence of overexpressed α-tubulin, we still saw similar results as in Fig 7: control comet speed (0.57 ± 0.12 μm/sec) was significantly slower than in GIP-1^late gut(-)^ (0.69 ± 0.12 μm/sec, p = 0.008 two-tailed t-test).

## Acknowledgements

We thank Jeremy Nance, Dan Dickinson, Bob Goldstein, Geraldine Seydoux, and Karen Oegema for providing strains, plasmids, protocols, and antibodies. We also thank members of the Feldman lab for helpful discussions about the manuscript. Some of the nematode strains used in this work were provided by the *Caenorhabditis* Genetics Center (CGC), which is funded by the NIH Office of Research Infrastructure Programs (P40 OD010440). This work was supported by a March of Dimes Basil O’Connor Starter Scholar Research Award and an NIH New Innovator Award DP2GM119136-01 awarded to J.L.F. M.D.S. is supported by the NIGMS NIH award F32GM120913-01.

## Author Contributions

J. Feldman, M. Sallee, T. Skokan, and J. Zonka designed the study, performed experiments, and analyzed data. J. Feldman, B. Raftrey, M. Sallee, T. Skokan, and J. Zonka made reagents and strains. J. Feldman, M. Sallee, and J. Zonka wrote the manuscript and made the figures.

## References

1. Pickett-Heaps JD (1969) The evolution of the mitotic apparatus: An attempt at comparative ultrastructural cytology in dividing plant cells. Cytobios 3: 257–280.

2. Baas PW, Deitch JS, Black MM, Banker GA (1988) Polarity orientation of microtubules in hippocampal neurons: uniformity in the axon and nonuniformity in the dendrite. Proc Natl Acad Sci U S A 85: 8335–8339.

3. Tassin AM, Maro B, Bornens M (1985) Fate of microtubule-organizing centers during myogenesis in vitro. J Cell Biol 100: 35–46.

4. Bacallao R, Antony C, Dotti C, Karsenti E, Stelzer EH, et al. (1989) The subcellular organization of Madin-Darby canine kidney cells during the formation of a polarized epithelium. J Cell Biol 109: 2817–2832.

5. Sanchez AD, Feldman JL (2016) Microtubule-organizing centers: from the centrosome to non-centrosomal sites. Curr Opin Cell Biol.

6. Feldman JL, Priess JR (2012) A role for the centrosome and PAR-3 in the hand-off of MTOC function during epithelial polarization. Curr Biol 22: 575–582.

7. Chabin-Brion K, Marceiller J, Perez F, Settegrana C, Drechou A, et al. (2001) The Golgi complex is a microtubule-organizing organelle. Mol Biol Cell 12: 2047–2060.

8. Efimov A, Kharitonov A, Efimova N, Loncarek J, Miller PM, et al. (2007) Asymmetric CLASP-dependent nucleation of noncentrosomal microtubules at the trans-Golgi network. Dev Cell 12: 917–930.

9. Wu J, Akhmanova A (2017) Microtubule-Organizing Centers. Annu Rev Cell Dev Biol 33: 51–75.

10. Paz J, Luders J (2017) Microtubule-Organizing Centers: Towards a Minimal Parts List. Trends Cell Biol.

11. Oakley BR, Oakley CE, Yoon Y, Jung MK (1990) Gamma-tubulin is a component of the spindle pole body that is essential for microtubule function in Aspergillus nidulans. Cell 61: 1289–1301.

12. Kollman JM, Polka JK, Zelter A, Davis TN, Agard DA (2010) Microtubule nucleating gamma-TuSC assembles structures with 13-fold microtubule-like symmetry. Nature 466: 879–882.

13. Kollman JM, Greenberg CH, Li S, Moritz M, Zelter A, et al. (2015) Ring closure activates yeast gammaTuRC for species-specific microtubule nucleation. Nat Struct Mol Biol 22: 132–137.

14. Janski N, Masoud K, Batzenschlager M, Herzog E, Evrard JL, et al. (2012) The GCP3-interacting proteins GIP1 and GIP2 are required for gamma-tubulin complex protein localization, spindle integrity, and chromosomal stability. Plant Cell 24: 1171–1187.

15. Zheng Y, Wong ML, Alberts B, Mitchison T (1995) Nucleation of microtubule assembly by a gamma-tubulin-containing ring complex. Nature 378: 578–583.

16. Wiese C, Zheng Y (2000) A new function for the gamma-tubulin ring complex as a microtubule minus-end cap. Nat Cell Biol 2: 358–364.

17. Hannak E, Oegema K, Kirkham M, Gönczy P, Habermann B, et al. (2002) The kinetically dominant assembly pathway for centrosomal asters in Caenorhabditis elegans is gamma-tubulin dependent. The Journal of Cell Biology 157: 591–602.

18. Srayko M, Kaya A, Stamford J, Hyman AA (2005) Identification and characterization of factors required for microtubule growth and nucleation in the early C. elegans embryo. Dev Cell 9: 223–236.

19. Rogers GC, Rusan NM, Peifer M, Rogers SL (2008) A multicomponent assembly pathway contributes to the formation of acentrosomal microtubule arrays in interphase Drosophila cells. Mol Biol Cell 19: 3163–3178.

20. Motegi F, Velarde NV, Piano F, Sugimoto A (2006) Two phases of astral microtubule activity during cytokinesis in C. elegans embryos. Developmental Cell 10: 509–520.

21. Toya M, Terasawa M, Nagata K, Iida Y, Sugimoto A (2011) A kinase-independent role for Aurora A in the assembly of mitotic spindle microtubules in Caenorhabditis elegans embryos. Nat Cell Biol 13: 708–714.

22. Sulston JE, Schierenberg E, White JG, Thomson JN (1983) The embryonic cell lineage of the nematode Caenorhabditis elegans. Dev Biol 100: 64–119.

23. Leung B, Hermann GJ, Priess JR (1999) Organogenesis of the Caenorhabditis elegans intestine. Dev Biol 216: 114–134.

24. Yang R, Feldman JL (2015) SPD-2/CEP192 and CDK Are Limiting for Microtubule-Organizing Center Function at the Centrosome. Curr Biol 25: 1924–1931.

25. Wang S, Tang NH, Lara-Gonzalez P, Zhao Z, Cheerambathur DK, et al. (2017) A toolkit for GFP-mediated tissue-specific protein degradation in C. elegans. Development 144: 2694–2701.

26. Zhang L, Ward JD, Cheng Z, Dernburg AF (2015) The auxin-inducible degradation (AID) system enables versatile conditional protein depletion in C. elegans. Development 142: 4374–4384.

27. DeRenzo C, Reese KJ, Seydoux G (2003) Exclusion of germ plasm proteins from somatic lineages by cullin-dependent degradation. Nature 424: 685–689.

28. Nance J, Munro EM, Priess JR (2003) C. elegans PAR-3 and PAR-6 are required for apicobasal asymmetries associated with cell adhesion and gastrulation. Development 130: 5339–5350.

29. Armenti ST, Lohmer LL, Sherwood DR, Nance J (2014) Repurposing an endogenous degradation system for rapid and targeted depletion of C. elegans proteins. Development 141: 4640–4647.

30. Edgar LG, McGhee JD (1988) DNA synthesis and the control of embryonic gene expression in C. elegans. Cell 53: 589–599.

31. Schumacher JM, Ashcroft N, Donovan PJ, Golden A (1998) A highly conserved centrosomal kinase, AIR-1, is required for accurate cell cycle progression and segregation of developmental factors in Caenorhabditis elegans embryos. Development 125: 4391–4402.

32. Hannak E, Kirkham M, Hyman AA, Oegema K (2001) Aurora-A kinase is required for centrosome maturation in Caenorhabditis elegans. The Journal of Cell Biology 155: 1109–1116.

33. Achilleos A, Wehman AM, Nance J (2010) PAR-3 mediates the initial clustering and apical localization of junction and polarity proteins during C. elegans intestinal epithelial cell polarization. Development 137: 1833–1842.

34. Nakamura M, Yagi N, Kato T, Fujita S, Kawashima N, et al. (2012) Arabidopsis GCP3-interacting protein 1/MOZART 1 is an integral component of the gamma-tubulin-containing microtubule nucleating complex. Plant J 71: 216–225.

35. Dhani DK, Goult BT, George GM, Rogerson DT, Bitton DA, et al. (2013) Mzt1/Tam4, a fission yeast MOZART1 homologue, is an essential component of the gamma-tubulin complex and directly interacts with GCP3(Alp6). Mol Biol Cell 24: 3337–3349.

36. Masuda H, Mori R, Yukawa M, Toda T (2013) Fission yeast MOZART1/Mzt1 is an essential gamma-tubulin complex component required for complex recruitment to the microtubule organizing center, but not its assembly. Mol Biol Cell 24: 2894–2906.

37. Hutchins JR, Toyoda Y, Hegemann B, Poser I, Heriche JK, et al. (2010) Systematic analysis of human protein complexes identifies chromosome segregation proteins. Science 328: 593–599.

38. Cota RR, Teixido-Travesa N, Ezquerra A, Eibes S, Lacasa C, et al. (2017) MZT1 regulates microtubule nucleation by linking gammaTuRC assembly to adapter-mediated targeting and activation. J Cell Sci 130: 406–419.

39. Strome S, Powers J, Dunn M, Reese K, Malone CJ, et al. (2001) Spindle dynamics and the role of gamma-tubulin in early Caenorhabditis elegans embryos. Mol Biol Cell 12: 1751–1764.

40. Goodwin SS, Vale RD (2010) Patronin regulates the microtubule network by protecting microtubule minus ends. Cell 143: 263–274.

41. Goshima G, Wollman R, Goodwin SS, Zhang N, Scholey JM, et al. (2007) Genes required for mitotic spindle assembly in Drosophila S2 cells. Science 316: 417–421.

42. Richardson CE, Spilker KA, Cueva JG, Perrino J, Goodman MB, et al. (2014) PTRN-1, a microtubule minus end-binding CAMSAP homolog, promotes microtubule function in Caenorhabditis elegans neurons. Elife 3: e01498.

43. Baines AJ, Bignone PA, King MD, Maggs AM, Bennett PM, et al. (2009) The CKK domain (DUF1781) binds microtubules and defines the CAMSAP/ssp4 family of animal proteins. Mol Biol Evol 26: 2005–2014.

44. Bouckson-Castaing V, Moudjou M, Ferguson DJ, Mucklow S, Belkaid Y, et al. (1996) Molecular characterisation of ninein, a new coiled-coil protein of the centrosome. J Cell Sci 109 (Pt 1): 179–190.

45. Wang S, Wu D, Quintin S, Green RA, Cheerambathur DK, et al. (2015) NOCA-1 functions with gamma-tubulin and in parallel to Patronin to assemble non-centrosomal microtubule arrays in C. elegans. Elife 4: e08649.

46. Ren J, Wen L, Gao X, Jin C, Xue Y, et al. (2008) CSS-Palm 2.0: an updated software for palmitoylation sites prediction. Protein Eng Des Sel 21: 639–644.

47. Bellanger JM, Gönczy P (2003) TAC-1 and ZYG-9 form a complex that promotes microtubule assembly in C. elegans embryos. Curr Biol 13: 1488–1498.

48. Giet R, McLean D, Descamps S, Lee MJ, Raff JW, et al. (2002) Drosophila Aurora A kinase is required to localize D-TACC to centrosomes and to regulate astral microtubules. J Cell Biol 156: 437–451.

49. Berdnik D, Knoblich JA (2002) Drosophila Aurora-A is required for centrosome maturation and actin-dependent asymmetric protein localization during mitosis. Curr Biol 12: 640–647.

50. Tsai MY, Zheng Y (2005) Aurora A kinase-coated beads function as microtubule-organizing centers and enhance RanGTP-induced spindle assembly. Curr Biol 15: 2156–2163.

51. Le Bot N, Tsai MC, Andrews RK, Ahringer J (2003) TAC-1, a regulator of microtubule length in the C. elegans embryo. Curr Biol 13: 1499–1505.

52. Roostalu J, Cade NI, Surrey T (2015) Complementary activities of TPX2 and chTOG constitute an efficient importin-regulated microtubule nucleation module. Nat Cell Biol 17: 1422–1434.

53. Woodruff JB, Ferreira Gomes B, Widlund PO, Mahamid J, Honigmann A, et al. (2017) The Centrosome Is a Selective Condensate that Nucleates Microtubules by Concentrating Tubulin. Cell 169: 1066–1077 e1010.

54. Lacroix B, Letort G, Pitayu L, Salle J, Stefanutti M, et al. (2018) Microtubule Dynamics Scale with Cell Size to Set Spindle Length and Assembly Timing. Dev Cell 45: 496–511 e496.

55. Lin TC, Neuner A, Flemming D, Liu P, Chinen T, et al. (2016) MOZART1 and gamma-tubulin complex receptors are both required to turn gamma-TuSC into an active microtubule nucleation template. J Cell Biol 215: 823–840.

56. Lynch EM, Groocock LM, Borek WE, Sawin KE (2014) Activation of the gamma-tubulin complex by the Mto1/2 complex. Curr Biol 24: 896–903.

57. O’Toole E, Greenan G, Lange KI, Srayko M, Müller-Reichert T (2012) The Role of γ-Tubulin in Centrosomal Microtubule Organization. PLoS ONE 7: e29795.

58. Nashchekin D, Fernandes AR, St Johnston D (2016) Patronin/Shot Cortical Foci Assemble the Noncentrosomal Microtubule Array that Specifies the Drosophila Anterior-Posterior Axis. Dev Cell 38: 61–72.

59. Lingle WL, Salisbury JL (2010) Altered Centrosome Structure Is Associated with Abnormal Mitoses in Human Breast Tumors. The American Journal of Pathology 155: 1941–1951.

60. Salisbury JL, Lingle WL, White RA, Cordes LE, Barrett S (1999) Microtubule nucleating capacity of centrosomes in tissue sections. J Histochem Cytochem 47: 1265–1274.

61. Pihan GA, Purohit A, Wallace J, Malhotra R, Liotta L, et al. (2001) Centrosome defects can account for cellular and genetic changes that characterize prostate cancer progression. Cancer Res 61: 2212–2219.

62. Brenner S (1974) The genetics of Caenorhabditis elegans. Genetics 77: 71–94.

63. Paix A, Wang Y, Smith HE, Lee CY, Calidas D, et al. (2014) Scalable and versatile genome editing using linear DNAs with microhomology to Cas9 Sites in Caenorhabditis elegans. Genetics 198: 1347–1356.

64. Dickinson DJ, Pani AM, Heppert JK, Higgins CD, Goldstein B (2015) Streamlined Genome Engineering with a Self-Excising Drug Selection Cassette. Genetics 200: 1035–1049.

65. Furuta T, Baillie DL, Schumacher JM (2002) Caenorhabditis elegans Aurora A kinase AIR-1 is required for postembryonic cell divisions and germline development. Genesis 34: 244–250.

66. Shelton CA, Bowerman B (1996) Time-dependent responses to glp-1-mediated inductions in early C. elegans embryos. Development 122: 2043–2050.

67. Schindelin J, Arganda-Carreras I, Frise E, Kaynig V, Longair M, et al. (2012) Fiji: an open-source platform for biological-image analysis. Nature Methods 9: 676–682.

68. Schneider CA, Rasband WS, Eliceiri KW (2012) NIH Image to ImageJ: 25 years of image analysis. Nature Methods 9: 671–675.

69. Wickham H (2009) ggplot2: Elegant Graphics for Data Analysis. Ggplot2: Elegant Graphics for Data Analysis: 1–212.

